# DNA methylation memory of pancreatic acinar-ductal metaplasia transition state altering Kras-downstream PI3K and Rho GTPase signaling in the absence of Kras mutation

**DOI:** 10.1101/2024.10.26.620414

**Authors:** Emily K.W. Lo, Adrian Idrizi, Rakel Tryggvadottir, Weiqiang Zhou, Wenpin Hou, Hongkai Ji, Patrick Cahan, Andrew P. Feinberg

**Author notes:** Correspondence to: Patrick Cahan, 733 N Broadway, MRB 653, Baltimore, MD 21205, Andrew P. Feinberg, 855 N. Wolfe St. Rangos 570 Baltimore, MD 21205.

## Abstract

A critical area of recent cancer research is the emergence of transition states between normal and cancer that exhibit increased cell plasticity which underlies tumor cell heterogeneity. Pancreatic ductal adenocarcinoma (PDAC) can arise from the combination of a transition state termed acinar-to-ductal metaplasia (ADM) and a gain-of-function mutation in the proto-oncogene *KRAS*. During ADM, digestive enzyme-producing acinar cells acquire a transient ductal epithelium-like phenotype while maintaining their geographical acinar organization. One route of ADM initiation is the overexpression of the *Krüppel-like factor 4* gene (*KLF4*) in the absence of oncogenic driver mutations. Here, we asked to what extent cells acquire and retain an epigenetic memory of the ADM transition state in the absence of oncogene mutation. We identified differential DNA methylation at Kras-downstream *PI3K* and *Rho*/*Rac*/*Cdc42* GTPase pathway genes during ADM, as well as a corresponding gene expression increase in these pathways. Importantly, differential methylation persisted after gene expression returned to normal. Caerulein exposure, which induces widespread digestive system changes in addition to ADM, showed similar changes in DNA methylation in ADM cells. Regions of differential methylation were enriched for motifs of KLF and AP-1 family transcription factors, as were those of human pancreatic intraepithelial neoplasia (PanIN) samples, demonstrating the relevance of this epigenetic transition state memory in human carcinogenesis. Finally, single-cell spatial transcriptomics revealed that these ADM transition cells were enriched for PI3K pathway and AP1 family members, linking epigenetic memory to cancer cell plasticity even in the absence of oncogene mutation.

## Introduction

Pancreatitis is a major risk factor for the development of pancreatic ductal adenocarcinoma (PDAC), one of the deadliest malignancies in the United States^1,2^. PDAC develops in the exocrine compartment of the pancreas, which comprises acini, the cells responsible for synthesizing digestive enzymes, and ductal epithelial cells which transport digestive juices to the small intestine. A characteristic feature of pancreatitis is a transition state called acinar-to-ductal metaplasia (ADM), in which acinar cells acquire a transient ductal-like phenotype. ADM supports tissue regeneration and protection in response to injury and is normally followed by reversion to a normal acinar phenotype^3^.

The combination of ADM with an oncogenic mutation in the *KRAS* gene, the most commonly mutated gene in pancreatic cancer^4,5^, drives the formation of neoplastic precursor lesions^3^. Although ADM alone is insufficient to drive the formation of neoplastic lesions, recent cancer research has placed more emphasis on the investigation of such cellular transition states prior to driver gene mutation and how they may underly or enable cancer initiation^6,7^. For example, the links between esophageal adenocarcinoma and Barrett’s esophagus and between gastric adenocarcinoma and gastric metaplasia^8^ highlight the key role that cell fate transition states play in cancer development. The epigenetic basis of these transition states is of particular interest, as the epigenome facilitates cell identity and memory^9–12^. Furthermore, the epigenome is also of particular interest in pancreatic cancer, as epigenetic changes have already been shown to facilitate the primary-to-metastasis transition without the acquisition of additional driver mutations^13–15^.

The role of the epigenome remains elusive in establishing and maintaining the ADM transition state. More specifically, DNA methylation, which confers stable, long-term epigenetic memory via maintenance across cell divisions^10,16^, has not been comprehensively profiled in ADM. Here, we investigate the DNA methylation landscape and corresponding transcriptional landscape of the mutation-free ADM transition state and the subsequent reversion to a normal acinar phenotype. We ask to what extent cells retain a memory of the ADM transition state via DNA methylation, examine potential regulators of altered DNA methylation during ADM, and investigate the relationship of the ADM transition state to human pancreatic cancer precursor lesions. These findings may inform the mechanisms not only of heritable epigenetic memory of transition states and inflammation, but also of how epigenetic alterations presage or even elicit genetic mutations in cancer.

## Results

### ADM differentially methylated regions are enriched in Kras-downstream PI3K and Rho GTPase pathway genes

We first sought to generate a mouse model that targeted ADM induction to acinar cells in the absence of oncogenic driver mutations. To this end, we bred *Ptf1a-rtTA*; *TRE-KLF4* (AK) mice, which enabled inducible, acinar-specific overexpression of human *Krüppel-like factor 4* (*KLF4*) (Fig. 1A). Klf4 is a known player in injury and inflammation^17,18^ and has been found to be necessary and sufficient for ADM in mouse models of pancreatitis^19^. A two-day administration of doxycycline in AK mice induced key histological features of acute pancreatitis, including prominent structural changes and vacuolation in acinar cells by day 2 (D2) and increased stromal and immune cell presence by day 4 (D4), followed by recovery to a normal acinar phenotype by day 7 (D7), while normal ductal cells were unchanged during the same time course (Fig. 1B). We also examined changes in pancreas weight over the time course. We observed that AK animals exhibited a decreased pancreas weight as a percent of total bodyweight, which was corroborated by animal weights in a widely used model of pancreatitis induction by caerulein (Table S1, Extended Data Fig. 1A). Nevertheless, the normal histology of pancreata at D7 and the high expression of *KLF4* and ADM marker gene *Krt19* in D2 relative to normal pancreata quantified by qPCR validated not only temporally appropriate induction of *KLF4* but also corresponding upregulation of and recovery from an ADM-associated phenotype (Fig. 1B-C). Taken together, these results demonstrate that targeted overexpression of *KLF4* to acinar cells is sufficient to reproduce the physiological characteristics of ADM, and that ADM can be induced in a cell-intrinsic manner.

**Figure 1.**
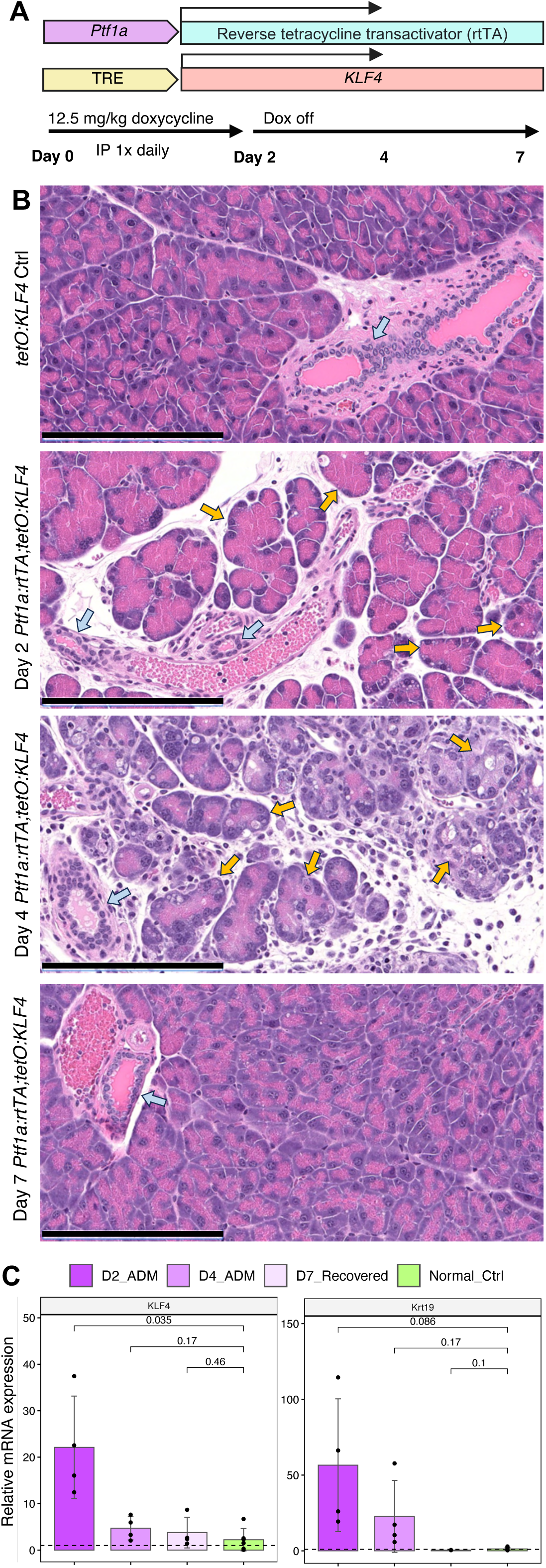
Genetically engineered mouse model of inducible acinar-specific KLF4 expression for induction of acinar-ductal metaplasia (ADM). A. Schematic of *Ptf1a-rtTA*, *TRE-KLF4* (AK) model and doxycycline administration schedule. TRE: Tet-response element. IP: Intraperitoneal. B. Representative hematoxylin and eosin staining of normal control, and Day 2, Day 4, and Day 7 pancreata as in administration schedule in (A). Blue arrows: normal ducts. Yellow arrows: ADM. Scale bar: 200 µm. C. Relative mRNA expression of murine Krt19 and human KLF4 as quantified by qRT-PCR. n = 2 mice. p-values: two-sided t-test. Error bars: SD.

To profile the DNA methylome of individual exocrine cell types during ADM, we performed whole-genome bisulfite sequencing (WGBS) on histologically pure tissue fragments of acini, ADM lesions, and ducts isolated using laser-capture microdissection (LCM) at days 2, 4, and 7 (Table S2, Extended Data Fig. 1B). Hundreds of microdissected regions were pooled per animal to achieve sufficient DNA yield and to provide robustness against variation that might result from profiling individual cuts. To validate the lack of canonical PDAC mutations in the AK model, we performed bisulfite-aware variant calling on all AK WGBS samples. We did not find any mutations in the top 4 genes mutated in PDAC (Kras, Tp53, Smad4, or Cdkn2a)^5,20^ in any sample (Table S3). Additionally, almost all genes with variants in the AK mice had an equivalent variant in the littermate control mice, suggesting that these are variants associated with the mouse strain and not associated with or induced by ADM.

Principal component analysis (PCA) on genome-wide CpG methylation grouped normal acini and ADM lesions separately, with recovered acini forming an intermediate group (Fig. 2A), demonstrating a detectable global difference in DNA methylation based on ADM status. When ducts were included, PCA grouped ducts separately from all other cell types at principal components 1 and 2 (Extended Data Fig. 1B), consistent with methylation studies in humans^21^. However, principal components 3 and 2 still demonstrated meaningful separation of normal acini and ADM lesions and an intermediate status of recovered acini (Extended Data Fig. 1C). Both genome-wide and at promoters specifically, there was no significant difference in CpG methylation level among normal acini, ADM lesions, and recovered acini, and methylation level was greater in ducts relative to the acinar-derived cell types (Extended Data Fig. 1C). However, examining DNA methylation globally often obscures important gene-level changes; thus, we next sought to identify specific genes with altered methylation during ADM.

**Figure 2.**
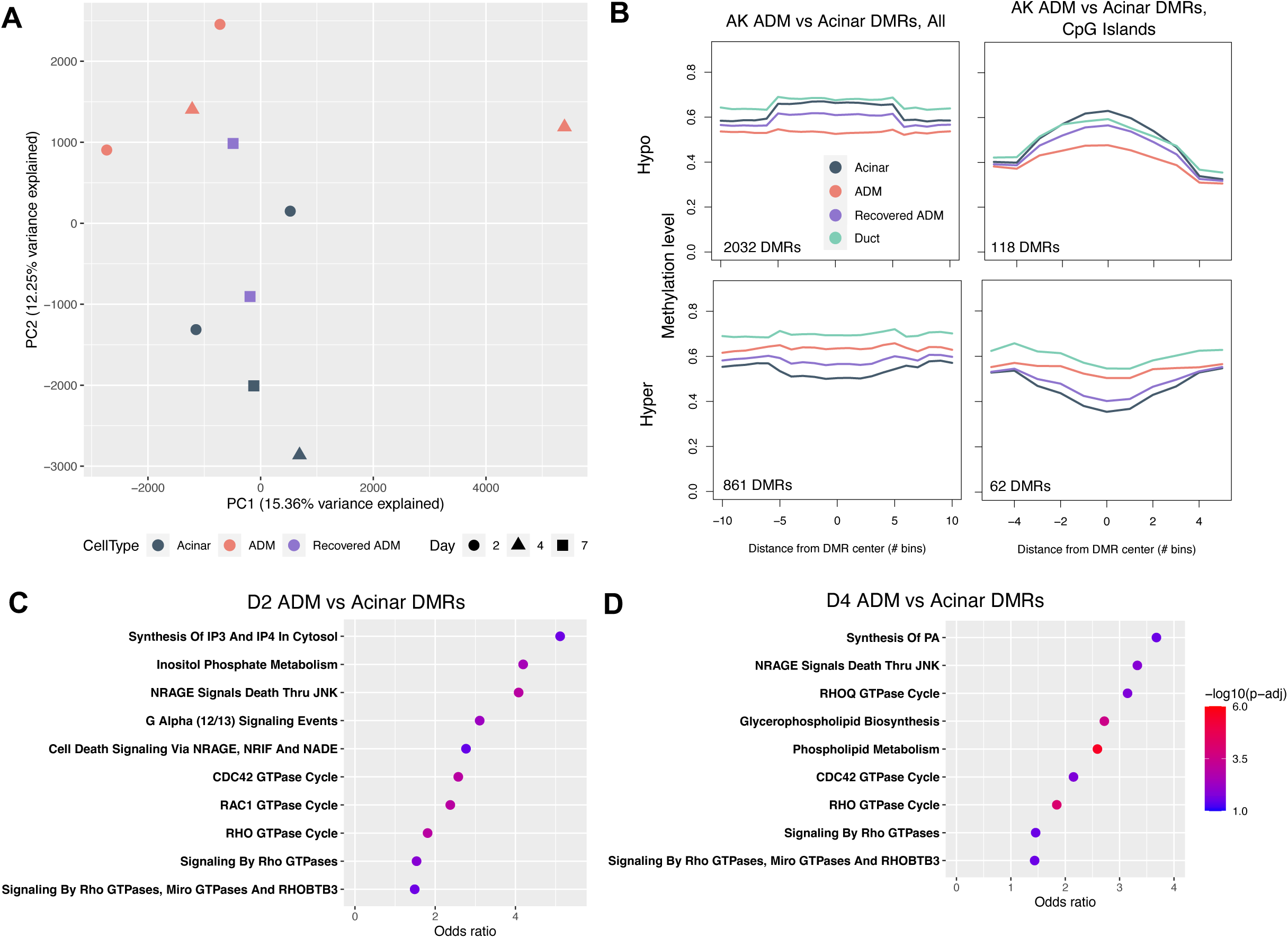
DNA methylation landscape of exocrine cell types during ADM. A. Principal component analysis (PCA) of laser capture microdissection-purified acini, ADM lesions, and recovered ADM lesions at PCs 1 and 2. B. Meta-region plots of AK ADM vs acinar DMRs summarized across all regions (left) and across CpG islands only (right) with 50% width buffer on each side. Methylation levels of recovered ADM lesions and ducts are overlayed. DMR: Differentially methylated region. C. Fisher’s gene set overrepresentation analysis (Fisher’s GSEA) of genes overlapping AK model D2 ADM vs acinar DMRs using Reactome 2022 Human gene sets. D. Fisher’s GSEA of genes overlapping AK model D4 ADM vs acinar DMRs using Reactome 2022 Human gene sets. Normal acinar, n = 4 mice; day 2 ADM, n = 2 mice; day 4 ADM, n = 2 mice; day 7 ADM, n = 2 mice. In B: duct, n = 10 mice.

We first performed mean-based differentially methylated region- (DMR-) finding to compare D2 and D4 ADM lesions to normal acinar controls in the AK model. Meta-region analysis of all samples at ADM vs acinar DMRs specifically demonstrated that D7 recovered samples, on average, had an intermediate methylation level between ADM and acinar samples, regardless of DMR directionality (Fig. 2B, Extended Data Fig. 2A). Analysis of genomic features overlapping these DMRs revealed an enrichment of DMRs over promoters, enhancers, and gene bodies, as well as CpG shores^22^ and shelves, relative to negative control randomly selected genomic regions (Extended Data Fig. 2B). Furthermore, Fisher’s exact gene set overrepresentation analysis (Fisher’s GSEA) of genes overlapping ADM vs acinar DMRs at both D2 and D4 (Table S4) revealed an enrichment of pancreatic acinar-related genes, reflecting an alteration of acinar identity during ADM (Extended Data Fig. 2C-D). Interestingly, Fisher’s GSEA also revealed a significant enrichment of genes related to inositol phosphate and phospholipid metabolism and signaling, including *Akt1*, several inositol phosphatases, and several phosphatidylinositol kinases (Fig. 2C-D, Extended Data Fig. 2E-F). *Akt1* is an essential node in the PI3K pathway, a well-known player in cell proliferation and oncogenesis, and enrichment of genes related to inositol phosphate signaling further suggest a major role of this pathway, in which a PI3K converts phosphatidylinositol-4, 5-bisphosphate (PIP2) to phosphatidylinositol-3, 4, 5-triphosphate (PIP3)^23^. In pancreatic carcinogenesis, members of the PI3K pathway are not often mutated; however, PI3K signaling is upregulated in human ADM, precursor lesions, and PDAC and can be inhibited to attenuate neoplastic growth^24,25^. We also observed the enrichment of genes related to Rho, Rac, and Cdc42 GTPase signaling, including many *Arhgef* and *Arhgap* family genes (Fig. 2C-D, Extended Data Fig. 2E-F). Rho/Rac/Cdc42 (R/R/C) GTPase pathways act in cytoskeletal organization, which is consistent with morphological cell remodeling observed in ADM. R/R/C GTPases also have known roles in cancer, particularly metastasis, due to their key role in regulating cell adhesion and migration^26,27^. Intriguingly, the PI3K and R/R/C pathways are downstream effectors of *Kras*, the most commonly mutated oncogene in PDAC^28^. We speculated that epigenetic changes during ADM in the Kras pathway, whose upregulation is a defining feature of pancreatic cancer, might reflect neoplastic-like activation of Kras even without a *Kras* mutation. Thus, we next examined to what extent cells undergoing ADM upregulate gene expression of Kras-driven pathways.

### DNA methylation changes in Kras downstream pathways are associated with gene expression changes

To determine whether the DNA methylation status of Kras-regulated pathways was reflected at the transcriptional level, we used 10X Visium spatial transcriptomics (ST) to profile gene expression in the same AK and caerulein pancreata on which LCM and WGBS were performed. Importantly, Visium ST allowed us to preserve the tissue architecture of our samples and minimize postmortem autolysis usually exacerbated by single-cell dissociation, while still profiling the samples in a whole-transcriptome manner. Despite the spot-level resolution, rather than single-cell resolution, of 10X Visium ST, the gene expression profiles of ST spots nevertheless generally clustered according to cell type, based on canonical marker gene expression (Fig. 3A, Extended Data Fig. 3A-C). To identify broad changes induced by KLF4 overexpression, initial differential gene expression analysis confirmed the recapitulation of ADM-associated signatures in our model. Specifically, 39 genes were upregulated at Day 2, including *Muc5ac* and *Tff1*; 836 genes were upregulated in Day 4, including *Reg3a*, *Reg3b*, *Reg3g*; and only 10 genes were higher in day 7 versus controls (Extended Data Fig. 4A-B).

**Figure 3.**
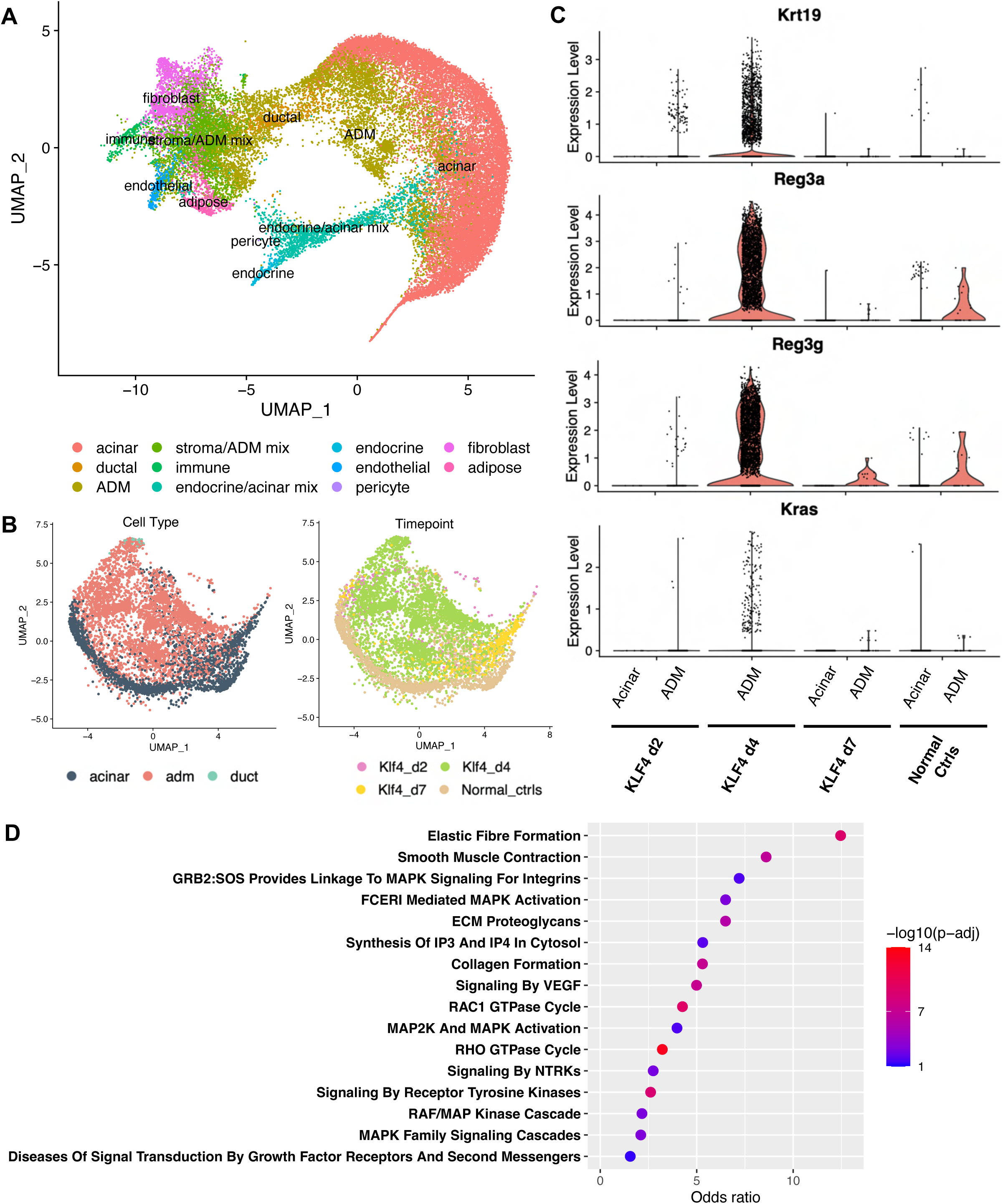
Gene expression patterns during AK ADM. A. UMAP projection of all AK mouse Visium ST spots (n = 31,620 spots). Cell types assigned via canonical marker gene expression. B. UMAP projection of AK mouse exocrine Visium ST spots only, labeled by cell type (left) and timepoint (right). C. Relative expression level of canonical ADM marker genes in acinar and/or ADM spots at each timepoint. D. Selected significantly enriched gene sets from Reactome 2022 Fisher’s GSEA performed on the 1412 genes significantly upregulated in highly pure ADM spots relative to highly pure acinar spots. Normal acinar, n = 4 mice; day 2 ADM, n = 2 mice; day 4 ADM, n = 2 mice; day 7 ADM, n = 2 mice.

To estimate the cell type composition of the ST spots, we trained the deconvolution method RCTD^29^ using a combination of “pseudo-spot” profiles derived from previously published single-cell data (Table S5) as well as from profiles of manually annotated histologically pure ST spots from a subset of the ST samples (Methods). Then, we applied RCTD to the remainder of the ST spots to assign cell type proportions to each spot and to identify highly pure exocrine spots (Fig. 3B). We noted that there were fewer ADM spots identified in the D2 sample; however, we attribute this to differences in the severity of the ADM phenotype at each time point and the area of the tissue section profiled by ST at each timepoint. D2 had a less severe histopathology compared to D4 in AK mice, and the tissue sections profiled for D2 were smaller in area than those profiled for D4; therefore, fewer ADM spots at D2 would be expected. The assigned spot identities corresponded well with the known timepoint annotations for each sample (Fig. 3B, Extended Data Fig. 3D). Increased expression of canonical ADM markers *Krt19*, *Reg3a*, and *Reg3g* in high-purity ADM spots relative to acinar spots further validated successful deconvolution by RCTD (Fig. 3C).

To examine if differential gene expression in ADM reflected differential methylation, we first compared highly pure ADM spots to highly pure acinar spots and identified more than 1400 significantly differentially expressed genes. Fisher’s GSEA of genes upregulated in ADM spots revealed enrichment of gene sets related to extracellular matrix and collagen formation; receptor tyrosine kinase signaling, including MAPK signaling; and Rho/Rac1 GTPase signaling (Fig. 3D). Examining the expression of specific genes overlapped by AK ADM vs acinar DMRs, many of the same PI3K and R/R/C GTPase pathway genes were indeed upregulated in ADM lesions, including PI3K-related *Akt1*, *Pip4k2b*, and *Ppard*, as well as R/R/C GTPase-related *Arhgef2*, *Gna13*, and *Rock2* (Fig. 4A-B). Fisher’s GSEA of simultaneously differentially methylated and differentially expressed genes confirmed joint enrichment of these pathways (Fig. 4C), demonstrating that pathway-level changes in gene expression during ADM corresponded well with DNA methylation changes. Another known pathway for activation of the Kras pathway in the absence of Kras mutations is via DNA methylation-associated silencing of RAS-effectors including RASSF1A^30^ or RASSF5 (NORE1A)^31^. We examined the methylation and gene expression status of these two genes, which were hypomethylated in D4 AK ADM relative to normal acini but were not differentially expressed. Taken together, these data support a model in which ADM cells specifically reprogram the PI3K and R/R/C pathways at both the transcriptional and DNA methylation levels, even in the absence of a Kras mutation.

**Figure 4.**
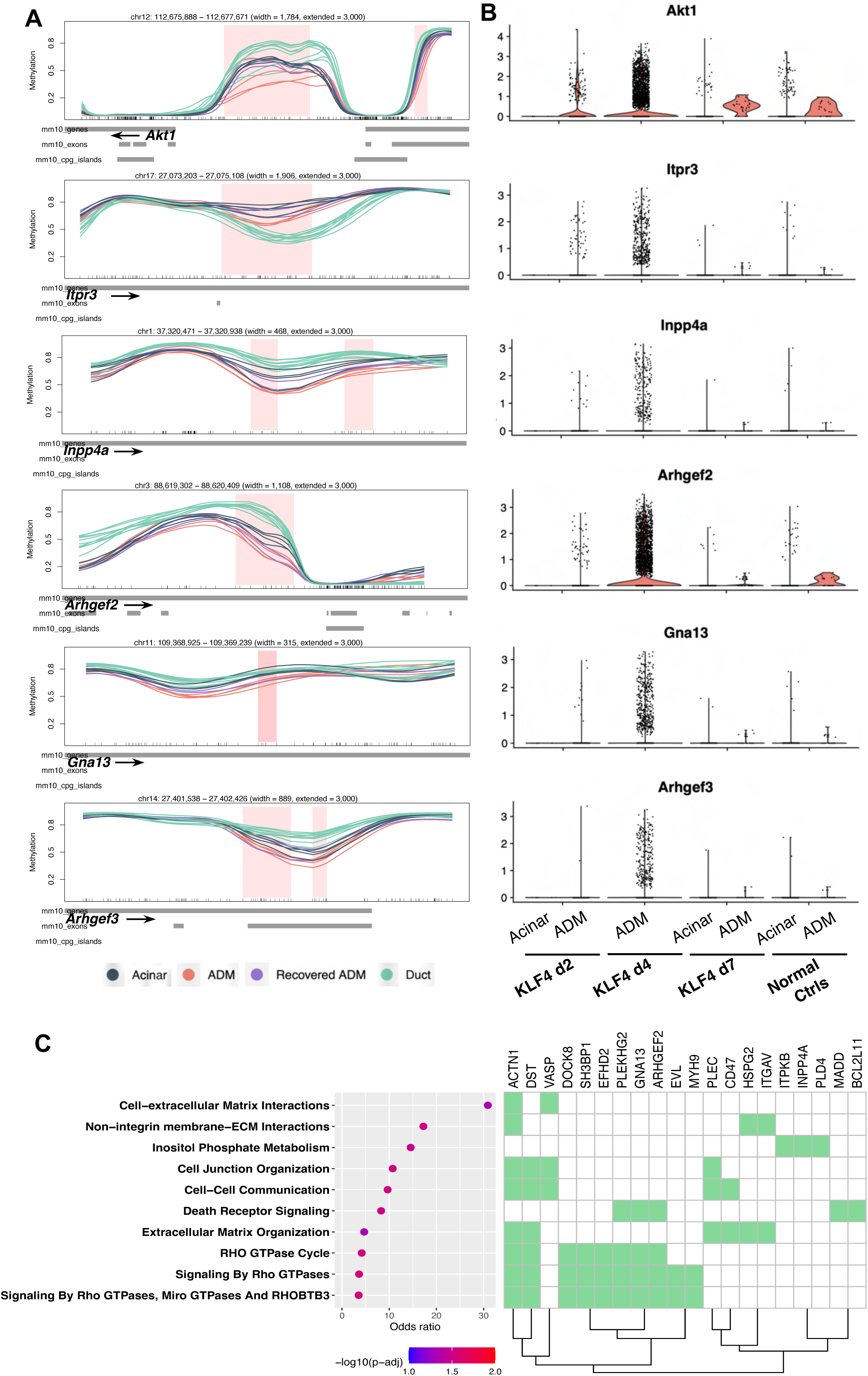
Integration of differential methylation and expression in AK ADM. Genomic line plots (A) and relative gene expression (B) for selected differentially methylated and differentially expressed PI3K pathway genes (*Akt1, Pip4k2b, Ppard*) and Rho GTPase family genes (*Arhgef2, Gna13, Rock2*) in ADM vs acini at each timepoint. In (A), tick marks on horizontal axis represent individual CpG sites. In (B), KLF4 d4 lacks an acinar violin plot as no spots in these samples were assigned an acinar identity during deconvolution. C. Reactome 2022 Fisher’s GSEA enrichment for genes both differentially methylated and differentially expressed in ADM vs acini. Normal acinar, n = 4 mice; day 2 ADM, n = 2 mice; day 4 ADM, n = 2 mice; day 7 ADM, n = 2 mice; duct, n = 10 mice.

### DNA methylation memory of the ADM transition state

To explore whether signatures of the ADM transition state are preserved even after resolution of ADM, we next asked whether DNA methylation changes seen during ADM were maintained in D7 acini. We identified regions of differential methylation comparing D7 acini to untreated acinar controls in the AK model. Meta-region analysis of all samples revealed that at D7 vs acinar DMRs, D7 acini, on average, had a more extreme methylation level than ADM lesions relative to normal acini, demonstrating that recovered acini did not simply have an intermediate methylation level between peak ADM and normal acini (Fig. 5A).

**Figure 5.**
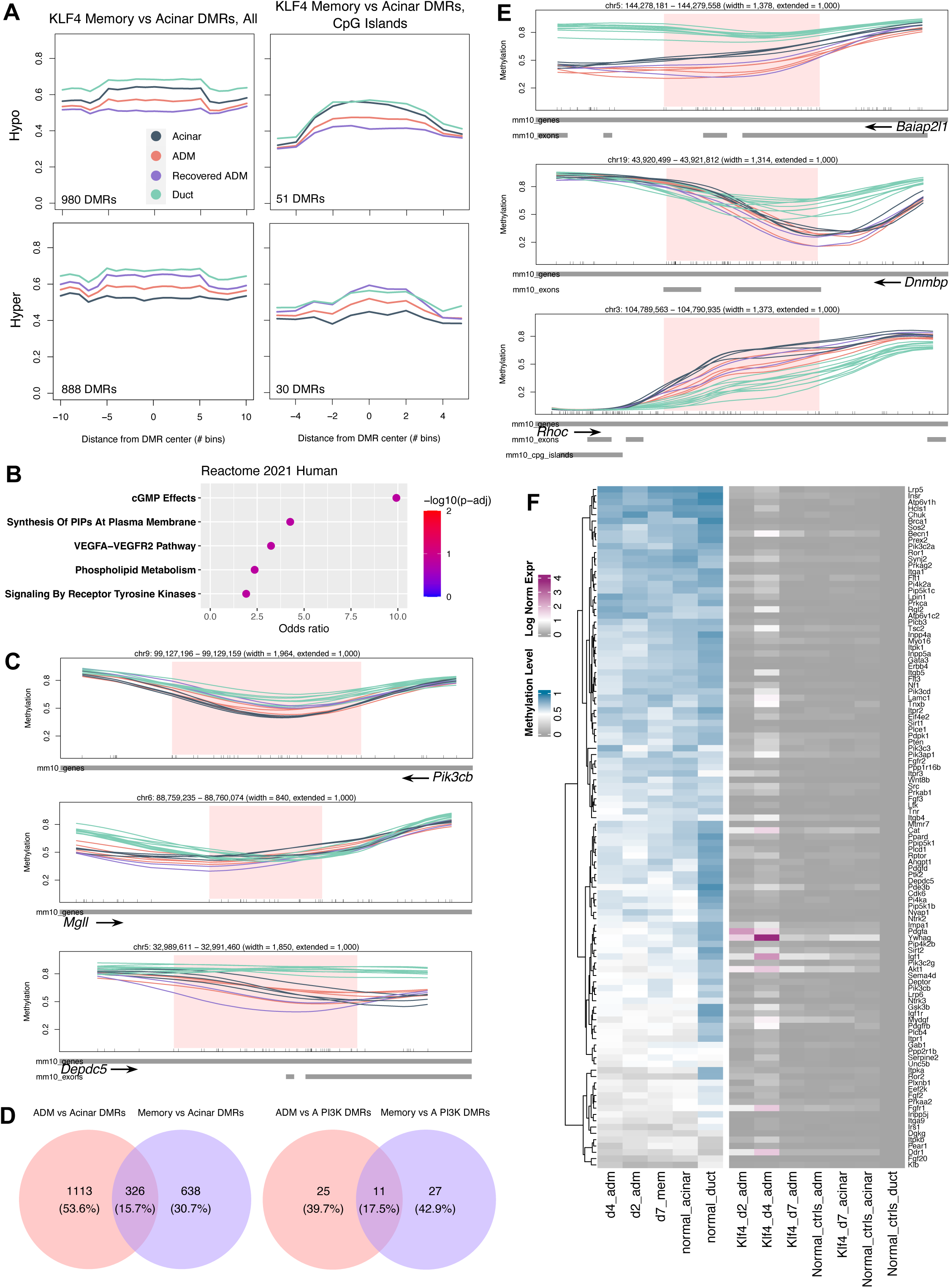
DNA methylation memory at PI3K pathway genes following ADM resolution. A. Meta-region plots of AK Day 7 (recovered ADM) vs acinar DMRs summarized across all regions (left) and across CpG islands only (right) with 50% width buffer on each side. B. Fisher’s GSEA of AK Day 7 (recovered ADM) vs acinar DMRs using Reactome 2022 Human gene sets. C. Genomic line plots of DNA methylation at representative PI3K-related genes. D. Overlap of ADM vs acinar DMRs and Day 7 vs acinar DMRs overlapping any gene (left) or overlapping PI3K pathway genes only (right). E. Genomic line plots of DNA methylation at representative Rho/Rac/Cdc42 GTPase-related genes. F. Summary heatmap of methylation level and normalized expression level at Reactome- and KEGG-annotated PI3K-related genes for each timepoint. In (C) and (E), tick marks on horizontal axes represent individual CpG sites.

To verify that these DMRs reflected memory of ADM DMRs at the same pathways, we performed Fisher’s GSEA on genes overlapping D7 vs acinar DMRs (Table S4). We observed a similar enrichment of PI3K pathway-related gene sets, including synthesis of PIPs at the plasma membrane and phospholipid metabolism (Fig. 5B). These genes included *Pik3cb*, *Tnxb*, and *Depdc5* (Fig. 5C, Extended Data Fig. 5A-B). We also determined that D7 vs acinar DMRs overlapped ADM vs acinar DMRs at 326 genes (or 15.7% of the total), indicating that the shared enrichment of PI3K represents methylation memory at many of the same genes (Fig. 5D). To exclude the possibility that the pathway enrichment we had observed in D7 vs acinar DMRs was driven by simply an intermediate methylation level relative to peak ADM during the process of recovery, we repeated Fisher’s GSEA on only DMRs where the D7 methylation level was more extreme than D2 and D4 ADM relative to normal acini. Enrichment of the PI3K pathway persisted, driven by genes including *Pten*, *Pik3cb*, and *Sos2* (Extended Data Fig. 5C). Although R/R/C GTPase-related genes were not significantly enriched at AK memory DMR genes, we still observed equal or as extreme D7 methylation in several R/R/C-related genes, including *Baiap2l1*, *Dnmbp*, and *Rhoc*, as in ADM (Fig. 5E).

To better assess the timeline of methylation memory and transcriptional resolution, we summarized the methylation and expression of known Reactome- and KEGG-annotated PI3K-related genes at each timepoint. We indeed observed that D7 expression was equivalent to normal control acinar expression, while D7 acini retained differential methylation relative to normal control acini (Fig. 5F). Systematic comparison of methylation and gene expression at known R/R/C-related genes showed a similar but not as drastic pattern (5Data Fig. 4D).

Combined, these results demonstrate that D7 recovered acini retain altered methylation, including at Pi3k pathway genes, even after expression returns to normal.

### AP-1, KLF, and pancreatic transcription factor binding is enriched at differentially methylated regions in ADM and in human PanIN lesions

We next investigated what TFs might be acting at ADM vs acinar DMRs to mediate changes in PI3K and R/R/C GTPase activity; thus, we performed TF motif enrichment analysis on these regions. As expected, the KLF4 motif was enriched in the AK D2 ADM vs acinar and D4 ADM vs acinar hypomethylated DMRs, as were those of KLF1 and KLF5, which was unsurprising given their almost identical reverse complementary motifs (Fig. 6A). Interestingly, we also observed an enrichment of AP-1-related motifs, including those of Jun and Fosl2, in D4 ADM vs acinar hypomethylated DMRs (Fig. 6A). Action of AP-1 family TFs including Jun and Fos is consistent with their known role as immediate early genes, or rapid responders to cellular stimuli, including inflammatory stress^33^, a characteristic induced by ADM. AP-1 family TFs also regulate a dynamic balance of cell proliferation and apoptosis and mediate oncogenic transformation, also consistent with processes of morphological and proliferative alterations that occur during ADM^34^.

**Figure 6.**
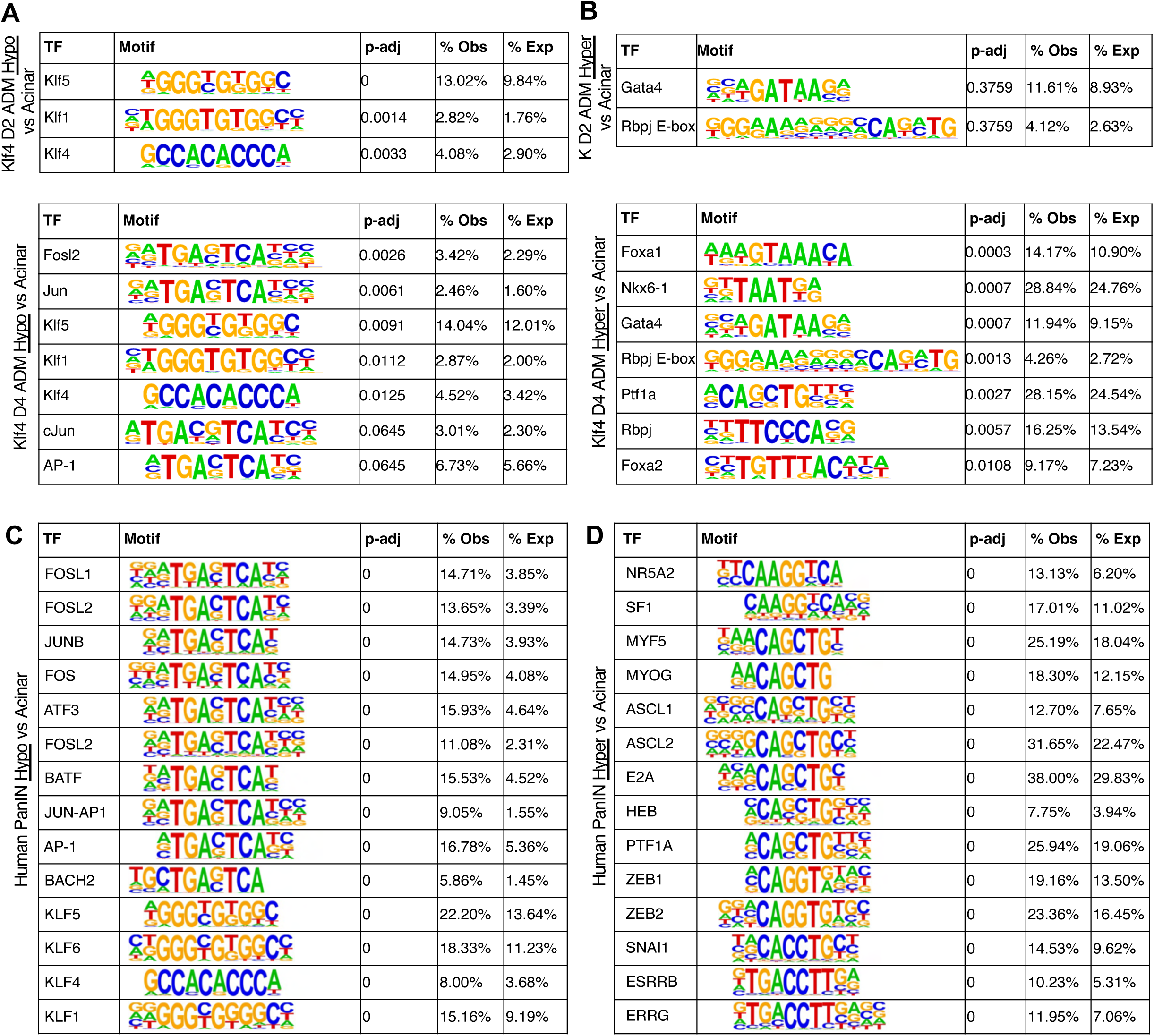
Consistent transcription factor motif enrichment at mouse ADM and human PanIN DMRs. A. Motif enrichment analysis results at AK ADM vs acinar hypo DMRs at Day 2 (top) and Day 4 (bottom). B. Motif enrichment analysis results at AK ADM vs acinar *hyper* DMRs at Day 2 (top) and Day 4 (bottom). A and B: normal acinar, n = 4 mice; day 2 ADM, n = 2 mice; day 4 ADM, n = 2 mice. Motif enrichment analysis results at human PanIN vs acinar hypo DMRs (C) and *hyper* DMRs (D). n = 8 acinar donors, n = 14 PanIN lesions from 8 donors. p-adj: Adjusted p-values via Benjamini–Hochberg correction.

When we investigated DMRs *hyper*methylated in D4 ADM relative to acini, we observed an enrichment of TF motifs related to pancreatic development, including those of Ptf1a, Rbpj, Foxa2, and Gata4 (Fig. 6B), potentially suggesting suppression of target genes that specify acinar identity during ADM. This is consistent with previous studies which suggest that loss of acinar identity promotes neoplasia and that maintenance of acinar identity is sufficient for neoplastic prevention. These prior studies show, for instance, that loss of expression of key acinar TFs Bhlha15 (aka Mist1) or Nr5a2 accelerates pancreatic neoplasia^35,36^, and that sustained Ptf1a overexpression prevents the formation of both precursor and malignant pancreatic lesions and can even revert neoplastic lesions to normal acini^37,38^. However, further studies will be needed to validate to what extent suppression of these key pancreatic TFs via epigenetic regulation can prevent or reverse neoplasia.

We then asked whether the observed TF enrichments reflected those of neoplastic pancreatic lesions in humans. We used DNA methylation data from available human acinar, ductal, and pancreatic intraepithelial neoplasia (PanIN) samples from our previous study that suggested that PanINs, pancreatic cancer precursor lesions, have an intermediate acinar-ductal DNA methylation signature at the gene level^21^. We examined known human PanIN vs acinar DMRs, again split into PanIN hypo- and hypermethylation. TF motif analysis on DMRs hypomethylated in PanIN lesions remarkably revealed the same enrichment of both AP-1 and KLF family motifs as in DMRs hypomethylated in ADM relative to acini (Fig. 6C, Table S6).

Furthermore, DMRs hypermethylated in PanIN lesions were enriched in motifs of TFs involved in pancreatic development, including NR5A2 and PTF1A, as well as in mesenchymal TFs ZEB1 and SNAI1, which had similar motif sequences to PTF1A (Fig. 6D). To assess the similarity of mouse and human TF enrichment in a more genome-wide manner, we compared the enrichment of all mouse ADM and human PanIN TF motifs. Generally, TF motifs highly enriched in mouse ADM vs acinar DMRs were also highly enriched in human PanIN vs acinar DMRs, and AP-1 family TFs and pancreatic development TFs were among the most highly enriched for mouse and human hypo- and hyper-methylated DMRs, respectively (Extended Data Fig. 6A-B). Taking further advantage of this human data, we also compared the genes overlapping human PanIN vs acinar DMRs themselves to those overlapping mouse ADM vs acinar DMRs. Fisher’s GSEA on these shared DMR genes identified the Rho GTPase pathway, which included several Arhgaps and Cdc42 GTPases in the overlapping set (Extended Data Fig. 6C). Together, these results suggest that the signatures and regulatory mechanisms of PanIN development may already be active at the pre-mutant ADM stage and do not require a Kras mutation.

### Epigenomic reprogramming of PI3K and R/R/C pathways in caerulein-induced ADM

To help corroborate our findings, we repeated the aforementioned histological and genomic assays using the widely used caerulein model of acute pancreatitis. Although caerulein administration induces ADM, it also has widespread inflammatory digestive effects including increased gastric secretion, gall bladder stimulation, and smooth muscle contraction in the gastrointestinal tract^39^, making it difficult to distinguish whether molecular features of ADM result from cell intrinsic activity as opposed to stromal, inflammatory, or systemic physiological influence. Nevertheless, we reasoned that the epigenetic and transcriptional features of ADM would still emerge, even in the presence of non-cell-intrinsic processes. Using the same administration timeline as in the AK model, we similarly observed acinar vacuolation and immune and stromal cell presence at peak pancreatitis on D2, formation of duct-like structures at D4, and resolution to a normal acinar phenotype by D7 (Extended Data Fig. 7A), consistent with the morphological features of AK model ADM.

CpG methylation profiles of cell types isolated using laser-capture microdissection (Table S2), as with AK cell types, grouped by ADM status via PCA (Extended Data Fig. 7B-D) even though global methylation levels among acinar-derived cell types did not differ significantly (Extended Data Fig. 7E). Combined PCA on ADM lesions, recovered acini, and normal acini from both the AK and caerulein studies clearly separated normal acini from ADM lesions at PCs 2 and 3, with recovered acini again forming an intermediate group (Extended Data Fig. 7D). This also supports a biological similarity between the AK and caerulein models of ADM. Examining differential methylation among cell types, the patterns of mean methylation level for ADM vs acinar comparisons in the AK model were less marked but nevertheless held true: at caerulein ADM vs acinar DMRs, D7 recovered samples held an intermediate methylation level between ADM and acinar (Extended Data Fig. 7F). At the gene level, Fisher’s GSEA of genes overlapping D2 and D4 ADM vs acinar DMRs in the caerulein model further validated the enrichment of genes related to GTPase and inositol phosphate signaling (Extended Data Fig. 7G, Table S4). To corroborate the gene expression signatures observed in the AK model, we also performed 10X Visium ST on caerulein model pancreata (Table S2), as well as deconvolution with RCTD as before. ST spots clustered according to cell type based on canonical marker gene expression as in the AK model (Extended Data Fig. 8A-C).

To evaluate DNA methylation memory in the caerulein model, we again compared D7 recovered acini to normal control acini. We similarly observed that in these DMRs, D7 acini had a more extreme methylation level than ADM lesions did relative to normal control acini (Extended Data Fig. 9A). Although there was no significant enrichment of PI3K-related pathways in genes overlapping these DMRs as seen in the AK model, we did observe significant enrichment of R/R/C-related gene sets (Extended Data Fig. 9B-C), perhaps suggesting model-specific patterns of DNA methylation memory.

Finally, we performed motif enrichment analysis on caerulein ADM vs acinar DMRs to examine if the same transcriptional regulators would be implicated. We observed the same enrichment of AP-1 family member motifs, including those of Batf, Atf3, Jun, Fosl1, and Fosl2 (Extended Data Fig. 9D-F, Table S6). Interestingly, we did not observe KLF family TFs enriched in DMRs where ADM samples were hypomethylated relative to acinar samples. At DMRs hypermethylated in ADM samples, we again saw motif enrichment of pancreatic development TFs, including Ptf1a, Rbpj, Nkx6-1, Foxa1, and Foxa2, similarly suggesting a repression of pancreatic identity during ADM (Extended Data Fig. 9E-F, Table S6). Taken together, these results suggest that features of the caerulein model are consistent with a cell-intrinsic induction of ADM.

### Single-cell resolution spatial transcriptomics reveals co-expression of multiple cell type signatures in cells undergoing ADM

We next asked if individual cells representing an acinar-ductal transition state could be identified and visualized. We reasoned that single cells with simultaneous expression of acinar, ductal, and even PanIN markers would represent a snapshot of the transition state. Thus, we performed single-cell resolution ST using MERFISH (multiplexed error-robust fluorescence in situ hybridization) on AK D2, D7, and normal control samples (Fig. 7A, Extended Data Fig. 10A-B). Our 500-target gene MERFISH panel comprised a mixture of canonical marker genes, differentially methylated genes identified via WGBS, differentially expressed genes identified via 10X Visium ST, and TFs identified via differential motif enrichment analysis (Table S7). In D2 ADM samples, we indeed identified individual cells simultaneously expressing acinar markers (*Ptf1a*, *Gp2*, *Pnliprp2*), ductal markers (*Krt19*, *Tff1*), and PanIN markers (*Muc5ac*) (Fig. 7A). Interestingly, these cells also expressed PI3K- and AP-1-related genes, including *Akt1*, *Itpr3*, *Pip5k1c*, and *Jund*. Visualization of single cells with co-expression of acinar, ductal, PanIN, PI3K, and AP-1 marker genes suggests that these cells are actively undergoing an acinar-ductal cell fate transition.

**Figure 7.**
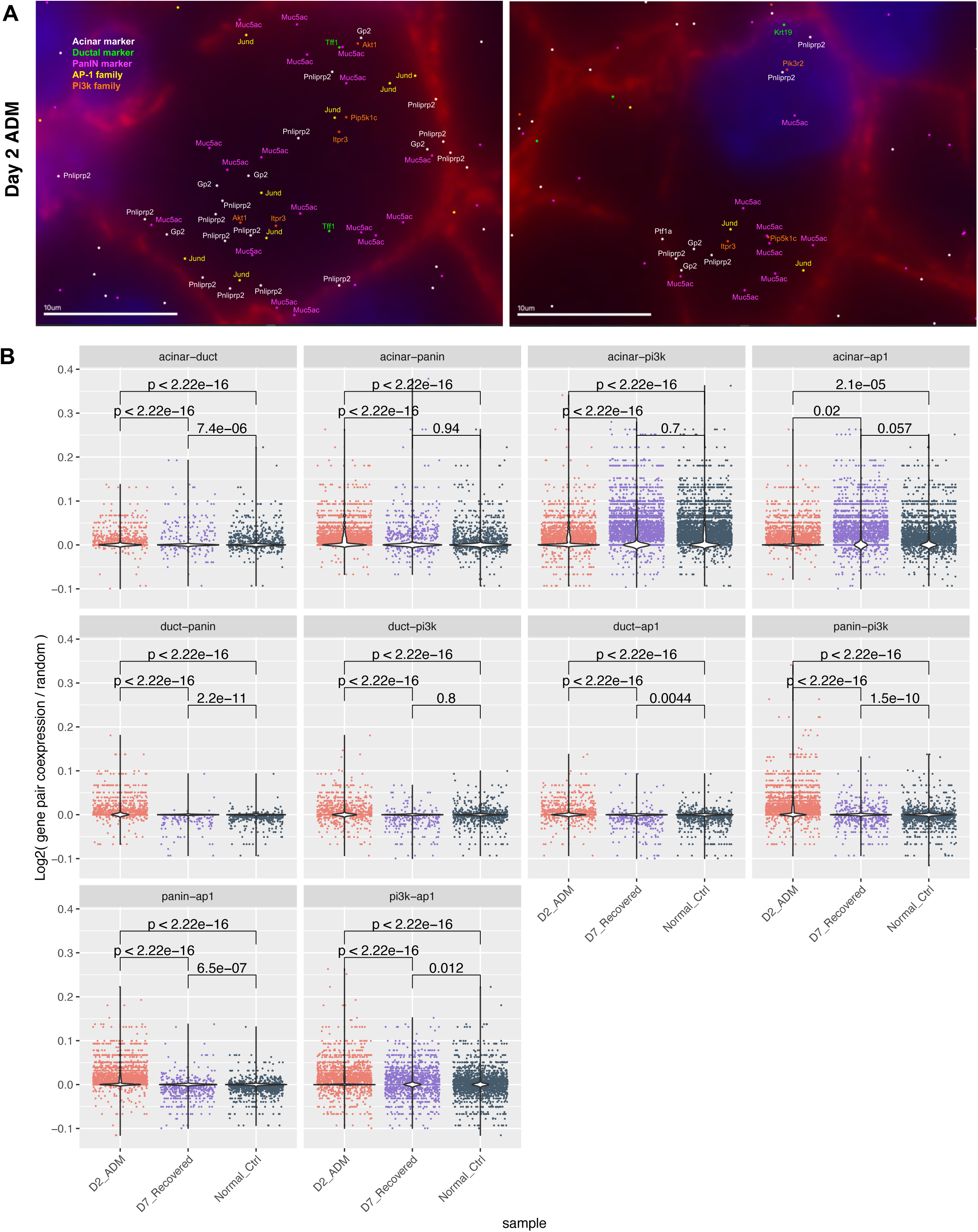
Joint gene expression signatures at a single-cell resolution. A. Example ADM cells co-expressing markers of acini (*Pnliprp2, Gp2*), ducts (*Tff1, Krt19*), panINs (*Muc5ac*), the PI3K pathway (*Akt1, Pip5k1c, Pik3r2, Itpr3*), and the AP1 complex (*Jund*). B. Violin plots of normalized gene pair co-expression proportion in each cell for the two labeled categories. n = 19776 cells. p-values: Wilcoxon rank sum test.

To quantify the abundance of cells in the acinar-ductal transition state among samples in a more systematic manner, we next examined pairwise co-expression of cell type signatures in all cells across all samples (Methods). We found that co-expression of acinar-duct, acinar-PanIN, and PanIN-duct marker gene pairs, normalized by total gene expression, total cell number, and random expectation, occurred most frequently in D2 ADM cells relative to D7 recovered and normal control cells (Fig. 7B), demonstrating increased abundance in D2 ADM of single cells actively in the acinar-ductal transition state. Interestingly, acinar-PI3K pathway co-expression and acinar-AP-1 family co-expression was less abundant in D2 ADM samples than in D7 recovered and normal control cells. We reasoned that this might be due to a decreased abundance of acinar markers in cells undergoing ADM and that ductal and PanIN markers would have more co-expression with PI3K and AP-1 family genes. Indeed, duct-PanIN, duct-PI3K, duct-AP-1, PanIN-PI3K, and PanIN-AP-1 marker co-expression was significantly more abundant in D2 ADM relative to D7 recovered and normal control cells. This further suggests that cells actively in the ADM transition state acquiring more duct-like characteristics also acquire PanIN-like characteristics and upregulate PI3K pathway genes and AP-1-related genes.

## Discussion

In this study, we examined epigenetic memory in acinar-ductal metaplasia, we investigated to what extent cells retain DNA methylation memory of the ADM state, and we evaluated the potential upstream regulators and downstream gene expression processes of DNA methylation changes in ADM. We found that during ADM, differential methylation was strongly induced at PI3K and R/R/C GTPase signaling genes, downstream effectors of Kras. Consistent with these changes, expression of genes in these pathways was also upregulated during peak ADM. However, differential methylation persisted at these genes following recovery, while gene expression returned to normal, demonstrating that recovered cells retain an epigenetic memory of the transition state at the level of DNA methylation. We implicated AP-1 family transcription factors as positive regulators of ADM, as their motifs were enriched at regions hypomethylated in ADM lesions relative to normal acini. We found that motifs of pancreatic development transcription factors were enriched at regions hypermethylated in ADM lesions. Additionally, we identified and visualized single cells actively in the ADM transition state, which simultaneously displayed duct- and PanIN-like characteristics and upregulated PI3K- and AP-1-related genes.

While we also used the caerulein model of acute pancreatitis to corroborate our findings, our study can attribute the observed changes in DNA methylation and gene expression to cell-intrinsic expression of KLF4. Finally, our study is highly relevant to human ADM, as the KLF4-induced ADM lesions observed here shared differentially methylated genes with human PanINs, and enrichment of AP-1 motifs and pancreatic development TF motifs was also consistent between mouse ADM and human PanINs. This shared epigenetic signature suggests that ADM lesions acquire characteristics of neoplasia even in the absence of oncogenic mutation. To further evaluate this claim, future studies profiling methylation in mouse models harboring Kras mutations, e.g. in the widely used Kras*^LSL-G12D/+^*; Pdx-1-Cre mouse model, will be necessary.

Though ours is the first to identify methylation reprogramming of the PI3K and R/R/C pathways in ADM, previous studies have also examined the behavior of PI3K and R/R/C in pancreatic cancer, although those studies also involved oncogenic mutations. PI3K signaling is almost universally activated in murine ADM, PanIN, and PDAC lesions, and inhibition or knockout of PI3K-downstream kinase PDK1 is known to prevent ADM, PanIN, and PDAC formation can be prevented *in vivo*^24^. Furthermore, the p110⍺ isoform of PI3K is known to regulate Rho and Rac1 GTPase activity in acinar cells, and inactivation of p110⍺ inhibits ADM and tumor formation *in vivo*^25,40^. There has also been emerging investigation of the relationship of PI3K signaling with cell plasticity. For example, homozygous H1047R gain-of-function mutation in *PIK3CA* has been found to impair the differentiation ability of human induced pluripotent stem cells and to increase their expression of stemness markers^41^. Additionally, human breast cancer samples display an association between increased PI3K signaling level and increased transcriptional cell stemness score^42^. Though further studies will be necessary to evaluate the extent to which the relationship between DNA methylation and transcriptional changes is causal, our study demonstrates DNA methylation-level reprogramming and memory of PI3K and R/R/C pathways in ADM in the absence of oncogenic mutation.

AP-1 also has other known roles in similar models of epithelial inflammation. In skin inflammation, sustained binding of JUN at chromatin domains following an inflammatory stimulus allows rapid re-recruitment of binding partner FOS in response to a secondary stimulus^43^. In mouse models of pancreatic inflammation and pancreatic cancer, AP-1 family motifs are enriched at regions of chromatin that are opened in response to inflammation and that remain open following oncogenic Kras mutation^7,44^. These independent findings are consistent with a role of AP-1 family members in epigenetic memory of ADM, though ours is the first to link AP-1 with DNA methylation alterations and memory in ADM. Future studies will help determine the mechanisms by which altered DNA methylation might directly or indirectly affect the behavior of AP-1 family members and pancreatic development TFs. For example, DNA methyltransferases are known to interact directly with AP-1 family members, including JUN, FOS, FOSL1, and FOSL2 *in vitro*^45^, providing a possible recruitment mechanism consistent with differential AP-1 binding at DMRs. Alternatively, TFs are known to preferentially bind motifs based on their methylation status^46,47^, which could also provide specificity to AP-1 and pancreatic TF binding patterns.

As novel methods are developed for performing simultaneous whole-transcriptome and whole-epigenome spatial profiling^48^, more will be revealed about the spatial and temporal dynamics of epigenetic memory in inflammation and metaplasia. Although methods for simultaneous transcriptome and DNA methylome profiling currently only exist at the dissociated single cell level^49^, already methods have been developed for spatial profiling of the transcriptome and chromatin accessibility simultaneously^50^, and for the transcriptome and histone modifications simultaneously^50,51^. In ADM, co-profiling of the transcriptome and DNA methylome in the same cell will allow precise detection of the point when transcription no longer reflects the methylation pattern of DNA, or in other words, the point when DNA methylation becomes a mark of memory rather than of active regulation.

We speculate that the sustained differential methylation of Kras-downstream pathways even after transcriptional recovery may reflect or even facilitate a selective pressure for sustained Kras activation via an oncogenic activating mutation. We theorize that this could be viewed as a type of oncogene addiction, a well-known characteristic of Kras-mutant cancers including PDAC^52^. Although oncogene addiction is classically viewed as a vulnerability of cancers harboring already-mutated oncogenes, our results could suggest that oncogene “addiction” may begin prior to mutation and may even create a selective pressure for mutation itself. The process of altered DNA methylation preceding oncogene upregulation, addiction, or mutation has precedents^53^. Consistent with this idea, *IDH*-mutant gliomas display DNA hypermethylation at insulator protein CTCF binding sites, resulting in upregulation, although not mutation, of proto-oncogene PDGFRA^54^. On the genome level, it is known that a consequence of global DNA hypomethylation, a known characteristic of most solid tumors^55,56^, is increased chromosomal instability and therefore increased proto-oncogene and tumor suppressor copy number aberrations^57,58^. Others have also hypothesized that epigenetic memory of a ductal-like state may be an evolutionary adaptation to more rapidly prevent cell injury in response to secondary inflammatory challenges^7,43^, which would also be consistent with our observations of DNA methylation memory. Further studies will inform the efficacy of potential chemopreventative measures that take advantage of pre-mutant oncogene addiction and epigenetic memory of ADM. The administration of a DNA methylation inhibitor such as azacytidine may be a useful tool for further investigation; however, because the directionality of DMRs between normal acini and ADM lesions is not always the same, unilateral inhibition of DNA methylation with a small molecule drug may not be the best approach for modulating the ADM phenotype. An approach that can specifically demethylate or methylate a targeted region might be a better suited approach for this, or perhaps the administration of Kras pathway inhibitors in at-risk populations prior to oncogenic Kras mutation.

More broadly, our study importantly provides a snapshot of the DNA methylome at key points during a cellular transition state associated with cancer initiation and opens the door to understanding underlying processes that may establish and maintain transition states. Our findings of persistent DNA methylation could have applications to cancers generally, especially those in which pre-neoplastic cellular transition states have not already been identified histologically, or might have application to identifying premalignant lesions through liquid biopsy.

## Methods

### Animal models

All animal procedures were performed in accordance with Johns Hopkins University Animal Care and Use Committee-approved protocols MO18M24 and MO21M20. *Ptf1a-rtTA*, *TRE-KLF4* (AK) mice were generated by crossing strain B6.129S6(SJL)-Ptf1atm^3.1(rtTA)Mgn^/Mmjax (Jackson Laboratories, cat. #036492-JAX) with strain FVB.Cg-Tg(tetO-KLF4)32831Rup/Mmjax (Jackson Laboratories, cat. #036730-JAX). Mice were genotyped using JAX-supplied primers and protocols. ADM was induced by administering 12.5 mg/kg doxycycline intraperitoneally once every 24 hours for 2 days. For caerulein-induced pancreatitis, C57BL/6J wild-type mice (Jackson Laboratories, cat. #000664) were administered 0.625 mg/kg caerulein (Millipore Sigma, cat. #C9026) intraperitoneally 4x daily for 2 days. All animals were between 6 and 28 weeks of age. Animals were euthanized according to AVMA guidelines prior to pancreatic extraction.

### Pancreatic extraction and processing

Following euthanasia, pancreata were immediately harvested. A 2-4 cm incision was made in the abdominal skin along the midline and then into the abdomen. Pancreata were exteriorized and removed using a combination of blunt and sharp dissection. The tail of each pancreas was immediately flash frozen in liquid nitrogen for 5 minutes, then transferred to −80°C for long-term storage. The head of each pancreas was placed in 10% neutral-buffered formalin for fixation for 24 hrs, then transferred to 70% ethanol. Tissue was then dehydrated in a graded ethanol and xylene series, then embedded in paraffin. Paraffin tissue blocks were stored long-term at 4°C.

### Histology

For hematoxylin and eosin (H&E) staining, formalin-fixed, paraffin-embedded (FFPE) tissue was sectioned at 4 µm thick onto charged slides (Fisherbrand #12-550-15). Tissue sections were rehydrated in xylenes and a graded ethanol series. Slides were incubated in Gill’s No. 2 hematoxylin for 1 minute, then blued in tap water for 5 min. Slides were then incubated in alcoholic eosin Y solution for 1 minute, then dehydrated in a graded ethanol and xylene series. Coverslips were mounted using Histomount mounting solution (ThermoFisher, cat. #008030).

### qRT-PCR

RNA was extracted from frozen pancreatic tissue using RNeasy Mini Kit (QIAGEN, # 74104) according to manufacturer’s instructions. RNA was reverse transcribed to cDNA according to manufacturer’s instructions for SuperScript IV Reverse Transcriptase (ThermoFisher #18090010). Assuming an 80% reverse transcription efficiency, qPCR was performed according to manufacturer’s instructions for iTaq Universal SYBR Green Supermix (BioRad #1725121). PCR and fluorescence quantification were performed using a BioRad CFX 384 Real-Time PCR System. Primer sequences for qPCR are available in Table S8.

### Laser capture microdissection (LCM)

Fresh frozen tissue samples were cryo-sectioned at 10 µm thick onto PEN membrane slides and stored at −20°C. Immediately prior to LCM, slides were fixed briefly in 70% ethanol, then stained with hematoxylin and eosin. LCM for the appropriate targeted cell type was performed using Zeiss PALM Microbeam on 10X or 20X magnification. The number of pooled cuts per cell type needed to reach a desired input of at least 20 ng DNA for WGBS varied for each sample. Genomic DNA was extracted using QIAamp DNA Micro Kit (Qiagen cat. #56304) according to manufacturer’s instructions for the “Isolation of Genomic DNA from Laser-Microdissected Tissues” protocol, with carrier RNA inclusion. To increase final DNA yield, samples were eluted twice, using the eluate from the first elution for the second, and incubating for 5 min each before centrifugation.

### Whole-genome bisulfite sequencing (WGBS)

Genomic DNA samples were quantified by Qubit dsDNA HS assay (ThermoFisher Scientific, cat. #Q32851). 1% unmethylated Lambda DNA (Promega cat. #D1521) was spiked into genomic DNA to monitor bisulfite conversion efficiency. xGen Methyl-Seq DNA Library Prep Kit (IDT, cat. #10009860) was used for WGBS library preparation. Genomic DNA (19-50 ng) was fragmented to a target peak of 300-400 bp using the Covaris S2 Focused-ultrasonicator in a 50 μl volume according to the manufacturer’s instructions. The fragmented genomic DNA was subjected to bisulfite conversion using the EZ DNA Methylation-Gold Kit (Zymo Research, cat. #D5005) following the manufacturer’s instructions. Methyl-seq libraries were generated according to the manufacturer’s instructions, and the resulting libraries were amplified for 7 PCR cycles using unique dual indexing primers (Swift Bioscience, cat. #X9096-PLATE). The resulting WGBS libraries were evaluated on a 2100 Bioanalyzer using the Agilent High-Sensitivity DNA Kit (Agilent Technologies, cat. #5067-4626) and quantified via qPCR using the KAPA Library Quantification Kit (KAPA Biosystems, cat. #KK4824). WGBS libraries were sequenced on an Illumina NovaSeq 6000 system at a 2 × 150 bp read length with a 5% spike-in of PhiX control library.

### WGBS variant calling

Bisulfite-aware variant calling was performed as previously described^21^. Briefly, python software Revelio was used to mask base positions potentially altered due to bisulfite treatment. Variant detector freebayes was used on Revelio-masked bam files to identify mutations in the coding sequences of the most commonly mutated genes in PDAC^5,20^. VCF (Variant Call Format) files outputted by freebayes were then converted into MAF (Mutation Annotation Format) files via vcf2maf. Finally, the Bioconductor package maftools was used to analyze the type and distribution of the given mutations, with a read coverage threshold of 30 and a variant detection threshold of 3 reads.

### WGBS analysis

Adapter sequences were computationally trimmed with Trim Galore (https://github.com/FelixKrueger/TrimGalore), with 10 additional bp trimmed from the 3’ end of R1 and 10 bp from the 5’ end of R2 to account for the low complexity polynucleotide tail added by the Adaptase enzyme. Bisulfite-aware alignment of the trimmed reads to the mm10genome was performed using Bismark with default parameters. Samtools was used to merge individual bam files corresponding to fastq file pairs and to name-sort the merged bam file. Bismark was then used to deduplicate merged, name-sorted bam files, then extract CpG methylation information. Bisulfite conversion rate was determined by alignment of trimmed reads to the lambda genome using Bismark with default parameters. Bisulfite conversion rate was at least 99.2% for all samples, with an average of 99.34% among all samples.

The Bioconductor package bsseq was used for integrating independent CpG_report files from Bismark’s methylation_extractor. CpG methylation values for each sample were smoothed with the BSmooth function using default parameters. Principal component analysis (PCA) was performed on smoothed methylation values generated by bsseq using the prcomp function from the R stats package, with parameters scale = TRUE and center = TRUE.

For differential methylation analysis, smoothed CpG methylation values from autosomes were coverage-filtered with a cutoff of 2X for each biological replicate. DMR-finding was limited to autosomes to help mitigate potential bias introduced by sex. For pairwise comparisons, BSmooth.tstat was used to compute t-statistics between groups, with k=21 and local.correct=TRUE. Significant DMRs were identified using t-statistic quantile cutoffs of 0.05 and 0.95, and these DMRs were filtered using a CpG number cutoff of n ≥ 5 and a mean methylation difference cutoff of 10%. If two CpGs within a DMR were more than 300bp apart, the DMR was split into two separate DMRs.

Fisher’s exact gene set enrichment analysis (Fisher’s GSEA) was performed using web program Enrichr, a gene set overrepresentation analysis software based on Fisher’s exact test. First, genes overlapping DMRs found by bsseq were identified. Genomic annotations for mm10 genes were generated via Bioconductor package annotatr, then DMR-gene intersections were identified using the findOverlaps function from Bioconductor package GenomicRanges. Gene lists were input into Enrichr, and output tables for Reactome 2022 enrichment were visualized using R package ggplot2.

### Visium FFPE spatial transcriptomics (ST)

Whole-transcriptome ST was performed using Visium Spatial Gene Expression Reagent Kits for FFPE (10X Genomics #1000185, #1000362, and #1000366). Mouse probe set Visium_Mouse_Transcriptome_Probe_Set_v1.0_mm10-2020-A was used. Prior to library preparation, the DV200 quality metric of RNA from FFPE pancreas tissue was first verified using RNA 6000 Nano Kit (Agilent #5067-1511) according to manufacturer’s instructions. All pancreas sections used as input for Visium ST had a DV200 of ≥ 57%, with most samples ≥70%. Library preparation was performed according to manufacturer’s instructions (CG000407, Rev. A), with the following custom parameters: 1) each H&E-stained section was imaged in 12 parts on a Nikon Eclipse Ti Research Photomicroscope at 4X magnification, then images were stitched post-capture using ImageJ; 2) 0.5 µL RNase Block (Agilent #300151) was added per sample (e.g. 2.2 uL for 4X + 10%) at pre-hybridization (Step 1.1.a) and probe hybridization (Step 1.1.g); and 3) PCR cycle number (Step 4.2.d) ranged from 14-17 depending on the Cq value from Step 4.1.e. Sample Index PCR (Step 4.2.a) for each sample was performed with a unique index well from Dual Index Plate TS Set A (10X Genomics #3000511). The 20 FFPE Visium libraries were sequenced on one lane of an S4 flowcell on an Illumina NovaSeq 6000 system at a 2 x 150 bp read length. Libraries were loaded onto the flowcell at a ratio to achieve at least 25,000 reads per spatial spot covered by tissue.

### MERFISH

MERFISH was performed using the Vizgen MERCSOPE platform. MERSCOPE sample verification (Vizgen #10400008) was performed on pancreas FFPE tissue to verify RNA and tissue quality according to manufacturer’s instructions (91600004, Rev D). MERSCOPE gene imaging (Vizgen #10400006) was performed on AK D2, D4, D7, and normal acinar samples according to manufacturer’s instructions (91600112, Rev B), with the following sample-specific parameters: 1) cell boundary stain 3 (Vizgen #10400009) was used, 2) non-resistant FFPE Tissue clearing protocol was performed, and 3) probe hybridization was performed for 48 hrs. Custom 500-gene panel (Vizgen #10400003, Panel ID 6c74b9d8-2814-4e73-97dd-435ea4062659, Serial Number CP1069) is detailed in Table S7.

### Analysis of Visium data

Visium data was preprocessed using spaceranger (v 1.3.0) against 10X Genomics-provided mm10 reference transcriptome build refdata-gex-mm10-2020-A to generate gene-by-spot expression matrices. If necessary, manual image alignment was performed using the 10X Loupe Browser (v 5.1.0). R package Seurat was used to log-normalize, integrate, and perform Leiden clustering on spot expression profiles.

Deconvolution tool RCTD was used to assign cell identity proportions to Visium spot expression data. RCTD was trained on a combination of manually annotated pure Visium spots and publicly available single-cell datasets. Manual annotation of spots with pure compositions of acini, adipose tissue, ADM, duct, or endocrine cells was performed using the 10X Loupe Browser. For supporting cell types including B cells, endothelial cells, fibroblasts, macrophages, pericytes, and T cells, five publicly available pancreatic single-cell datasets (Table S5) were normalized by SCTransform and integrated with reciprocal PCA using Seurat. To create training profiles that would be comparable to manually annotated spots in terms of order of magnitude of counts, pseudobulk profiles were created by summing the counts by gene of 10 randomly selected cells of the same type. RCTD was performed using the following parameters: gene_cutoff = 0.000125, fc_cutoff = 0.5, UMI_min = 100, UMI_max = 2E+07, UMI_min_sigma = 300, CELL_MIN_INSTANCE = 25, CONFIDENCE_THRESHOLD = 10, doublet_mode = "full". Following RCTD deconvolution, only exocrine spots in the top 90^th^ percentile of a given cell type proportion with no other greater cell type proportion were used for downstream differential expression analysis with Seurat.

### Analysis of MERFISH data

Raw MERSCOPE data was preprocessed using the MERSCOPE Visualizer. Processed Vizgen datasets were log-normalized and integrated with reciprocal PCA using Seurat. Cell type co-expression analysis was performed using a list of canonical marker genes for each cell type (Table S7). Genes from the target gene panel related to PI3K and AP-1 were annotated based on KEGG, GO BP, and Reactome gene sets from MSigDB. For each cell, to normalize for cell-cell variability in MERSCOPE transcript detection, the proportion of unique gene pair combinations of x number of markers of cell type A and y number of markers of cell type B out of the total possible unique gene pairs in a cell was calculated. To normalize for random expectation, for each cell we then calculated the co-expression probability of x and y randomly selected markers without replacement. We reported the log2 fold change (signal/random) for each cell in each sample per A-B cell type pair.

### Transcription factor motif analysis

Transcription factor motif analysis at DMRs was performed using software package Homer. For each differential methylation comparison, a .txt file containing specifying the genomic coordinates of each DMR, plus 50bp of buffer on each side, was generated in R. Homer’s findMotifsGenome.pl program was run on each .txt file with mm10 genome and -size 200.

## Availability of data and materials

Raw FASTQ files for WGBS and 10X Visium data will be publicly available in Gene Expression Omnibus (GEO) at accession GSE249143 following publication. All other raw data are available upon reasonable request to the corresponding authors.

## Author Contributions

EKWL, APF, and PC conceived and designed the project. EKWL, AI, and RT acquired the data. EKWL, WZ, WH, HJ, PC, and APF analyzed and interpreted the data. EKWL, PC, and APF drafted and revised the manuscript.

## Acknowledgments

We thank Genevieve Stein-O’Brien, Dimitri Sidiropoulos, Zachary Nicholas, and Clare Sengupta for access and training on MERSCOPE. We thank Johns Hopkins Research Animal Resources for assistance with mouse procedures and Johns Hopkins Oncology Tissue Services, Amy Smith, Heather Kulaga, for assistance with tissue and histological procedures. We thank David Tuveson and Ralph Hruban for advice during this work.

## Inventory of Supporting Information

**Extended Data Figures 1-10**

**Supplementary Tables S1-9**

**Extended Data Figure 1.**
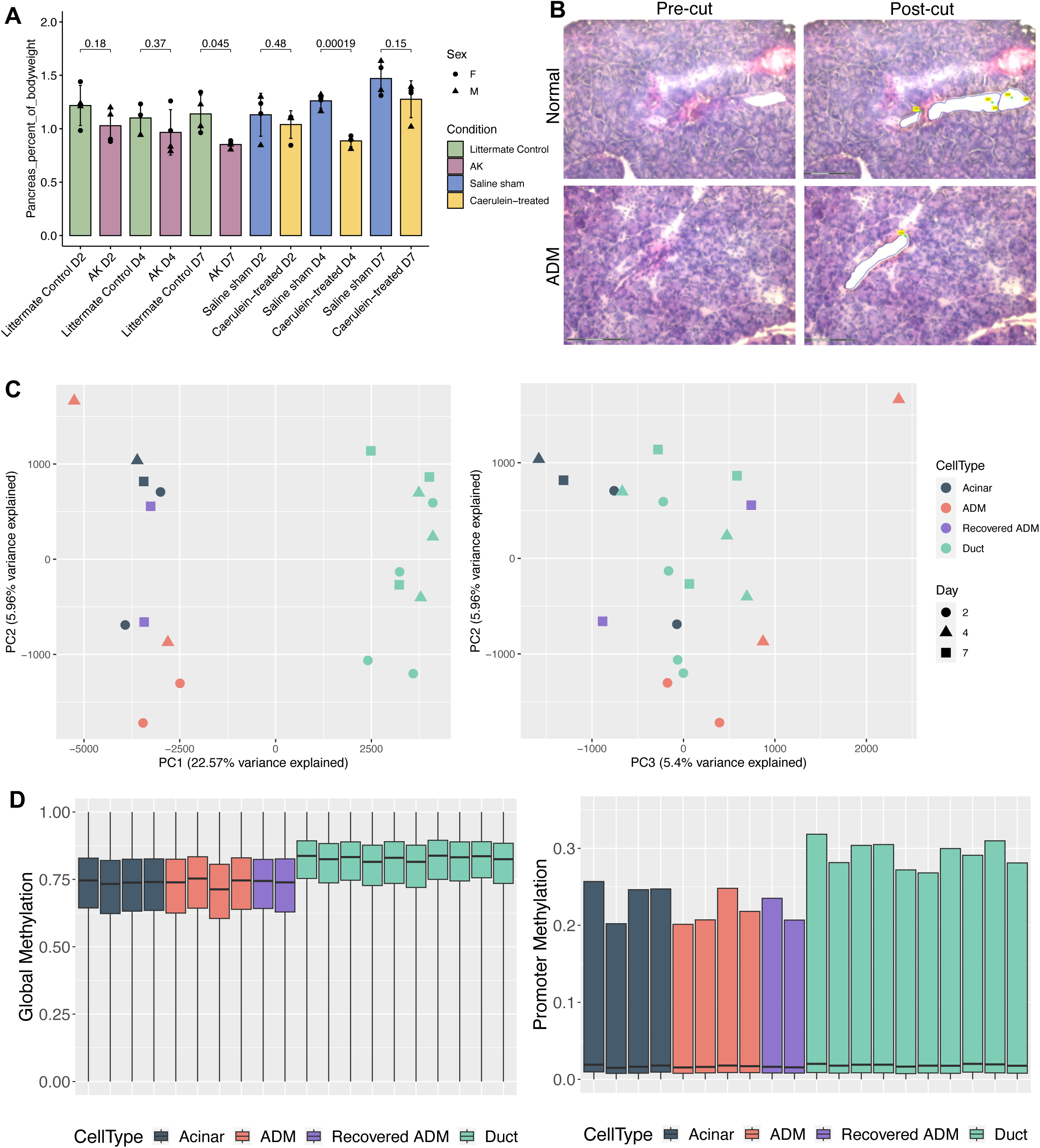
Laser-capture microdissection (LCM) and whole-genome bisulfite sequencing (WGBS) of exocrine cell types in mouse models of ADM. A. Representative examples of pre- and post-LCM cuts for ducts isolated from normal and ADM H&E-stained tissue. B. Principal component analysis (PCA) of genome-wide CpG methylation for all exocrine samples in AK model of ADM. C. Boxplots of genome-wide (left) and promoter-specific (right) CpG methylation values for all exocrine samples in AK model of ADM. Center line, median; box limits, upper and lower quartiles; whiskers, 1.5x interquartile range. Normal acinar, n = 4 mice; day 2 ADM, n = 2 mice; day 4 ADM, n = 2 mice; day 7 ADM, n = 2 mice; duct, n = 10 mice.

**Extended Data Figure 2.**
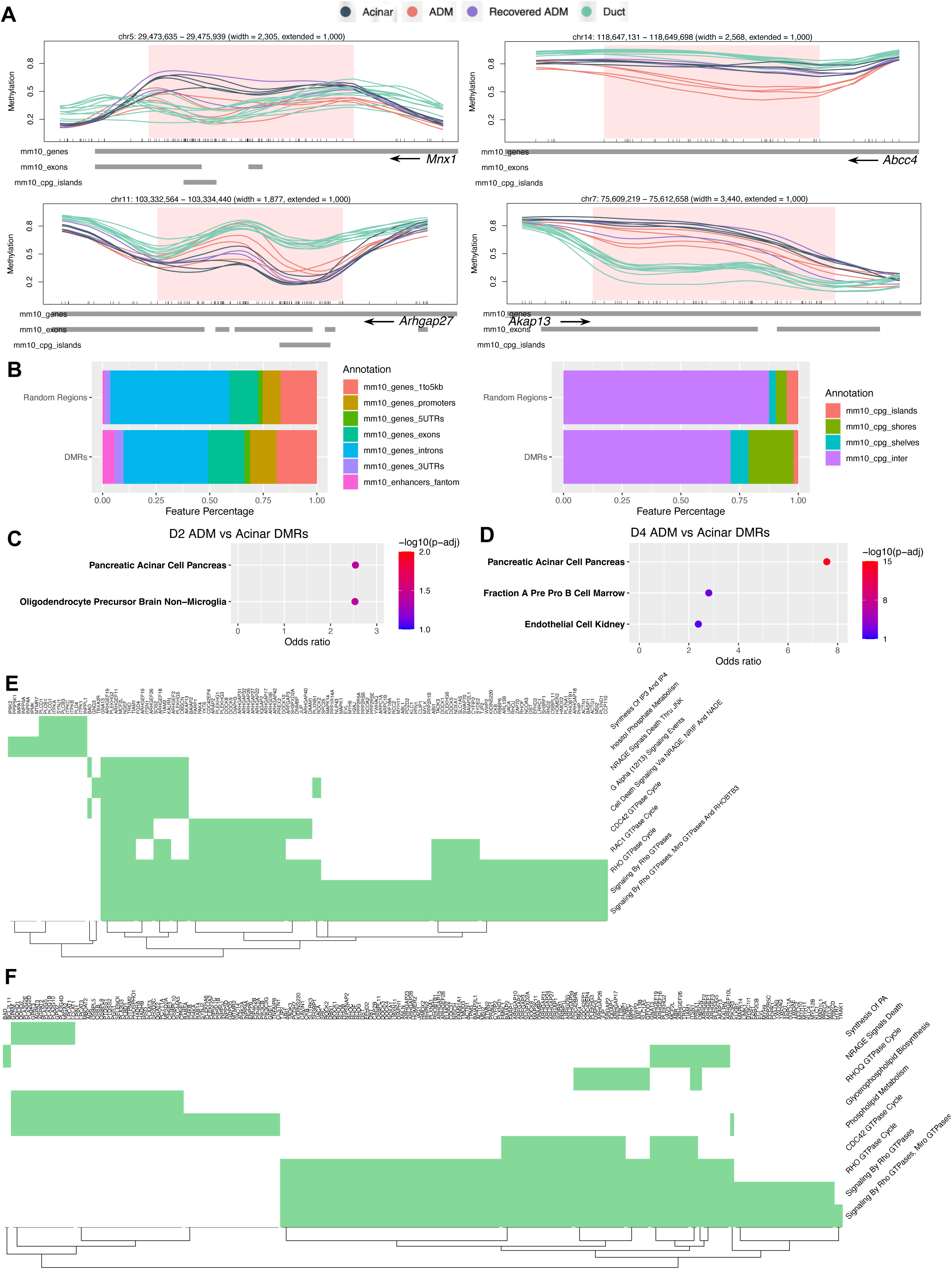
ADM vs acinar DMR profiles in AK ADM model. A. Genomic line plots of methylation level at representative DMRs comparing ADM lesions (D2 and D4) to normal acini. Tick marks on horizontal axes represent individual CpG sites. Normal acinar, n = 4 mice; day 2 ADM, n = 2 mice; day 4 ADM, n = 2 mice; day 7 ADM, n = 2 mice; duct, n = 10 mice. B. Stacked barplot of genomic region distribution of AK ADM vs acinar control DMRs as compared to randomly selected genomic regions of the same sizes. C. Fisher’s GSEA of genes overlapping AK model D2 ADM vs acinar DMRs using Tabula Muris gene sets. D. Fisher’s GSEA of genes overlapping AK model D4 ADM vs acinar DMRs using Tabula Muris gene sets. E. Genes (rows) overlapping AK D2 ADM vs normal acinar DMRs that contribute to each significantly enriched gene set (columns) as in Fig. 2C. F. Genes (columns) overlapping AK D4 ADM vs normal acinar DMRs that contribute to each significantly enriched gene set (rows) as in Fig. 2D.

**Extended Data Figure 3.**
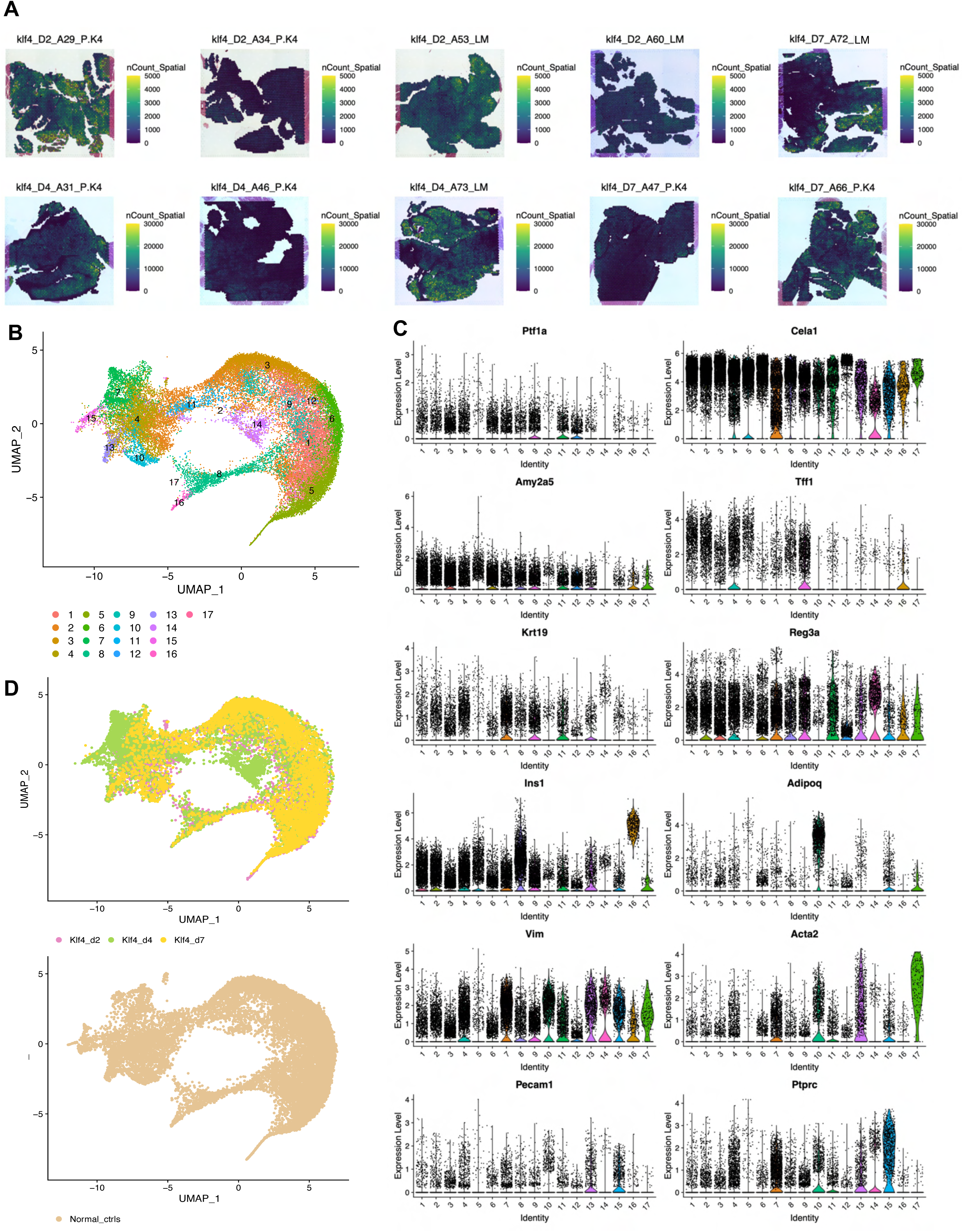
10X Visium Spatial Transcriptomics profiling. A. Per-spot count distribution for all AK model pancreata. Normal acinar, n = 4 mice; day 2 ADM, n = 2 mice; day 4 ADM, n = 2 mice; day 7 ADM, n = 2 mice; duct, n = 10 mice. B. UMAP projection of leiden clustering for all AK model ST spots (n = 31,620 spots). C. Normalized expression levels of canonical acinar (Ptf1a, Cela1, Amy2a5), duct (Tff1, Krt19), ADM (Reg3a), endocrine (Ins1), adipose (Adipo1), fibroblast (Vim), endothelial (Acta2, Pecam1), and immune (Ptprc) marker genes by cluster. D. UMAP projection of AK model spots by timepoint (top) and normal control spots (bottom).

**Extended Data Figure 4.**
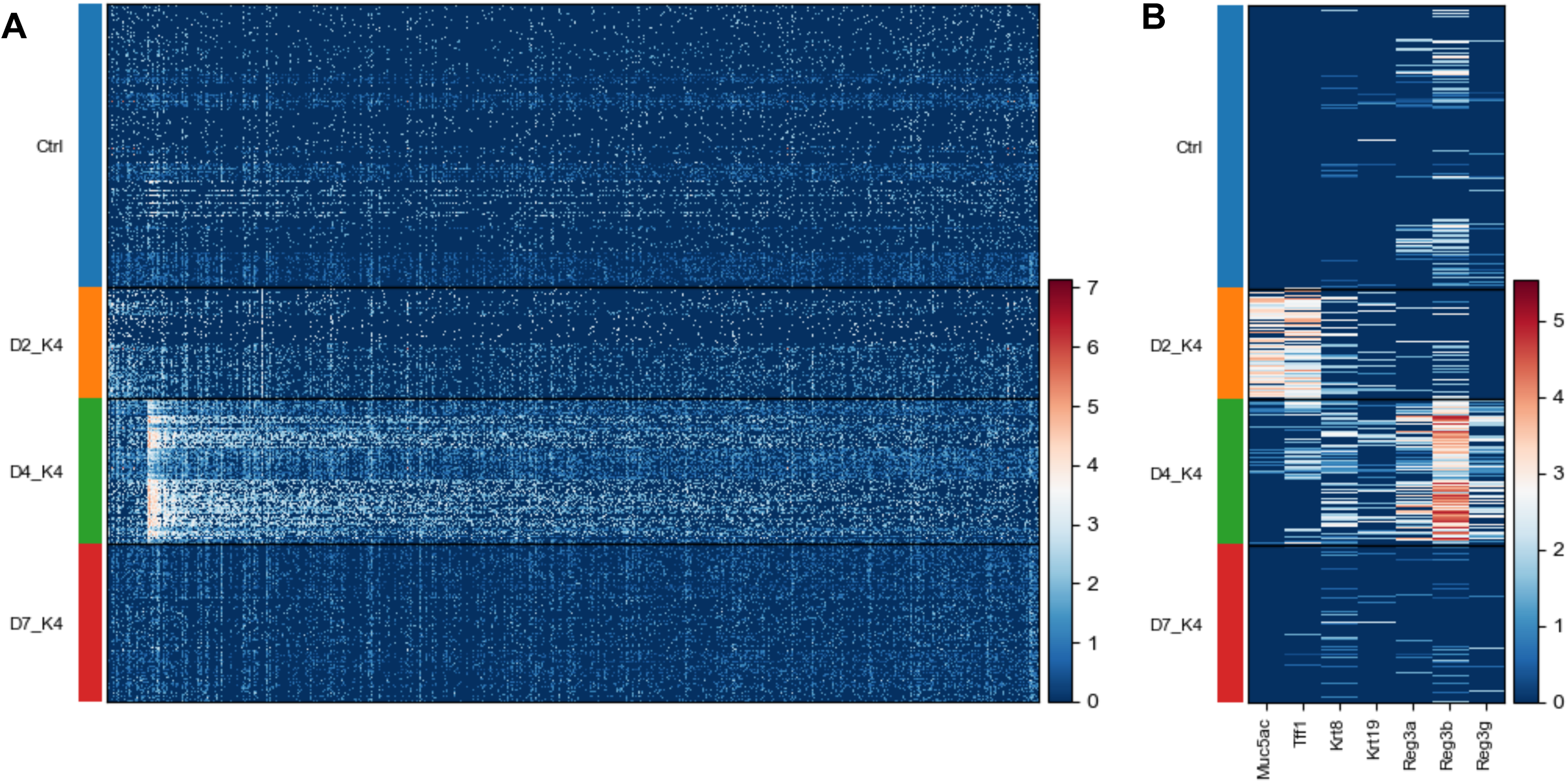
Gene expression differences in AK model as profiled by Spatial Transcriptomics. A. Relative gene expression of genes upregulated at D2, D4, and D7 in AK model versus controls. B. Relative gene expression of selected marker genes in AK model.

**Extended Data Figure 5.**
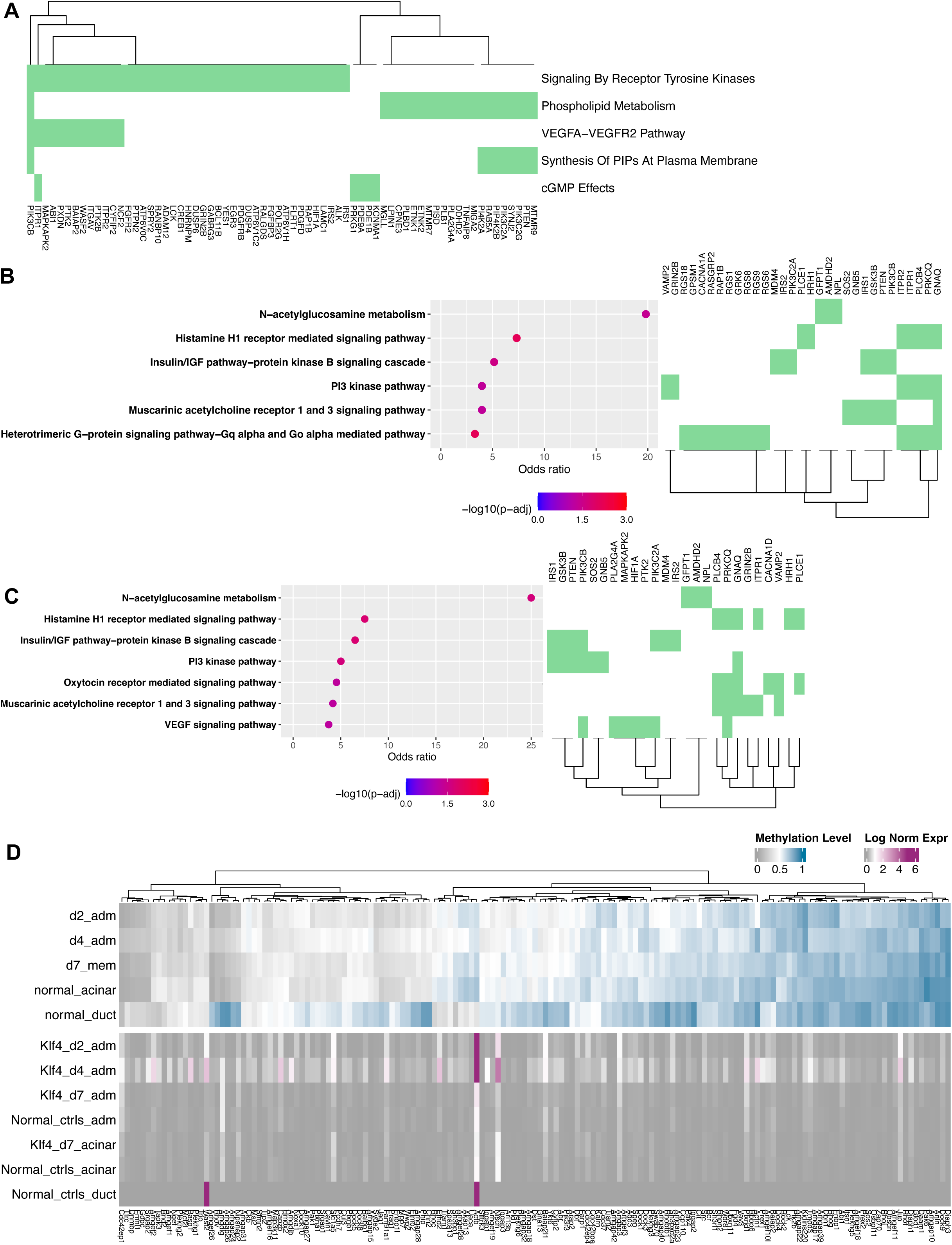
Memory DMR profiles for AK model pancreata. A. Genes (columns) overlapping AK model D7 vs normal acinar DMRs that contribute to each enriched Reactome gene set (rows) as in Fig. 5B. B. Fisher’s GSEA of genes overlapping AK model D7 vs normal acinar DMRs using Panther 2016 gene sets (left), with corresponding genes contributing to each gene set (right). C. Fisher’s GSEA of genes overlapping AK model D7 vs normal acinar DMRs for which the average D7 methylation value was more extreme than that of D2 & D4 ADM relative to normal acini controls. Fisher’s GSEA performed with Panther 2016 gene sets (left), with corresponding genes contributing to each gene set (right). D. Summary heatmap of methylation level and normalized expression level at Reactome- and KEGG-annotated Rho/Rac/Cdc42 GTPase-related genes for each timepoint. Normal acinar, n = 4 mice; day 2 ADM, n = 2 mice; day 4 ADM, n = 2 mice; day 7 ADM, n = 2 mice; duct, n = 10 mice.

**Extended Data Figure 6.**
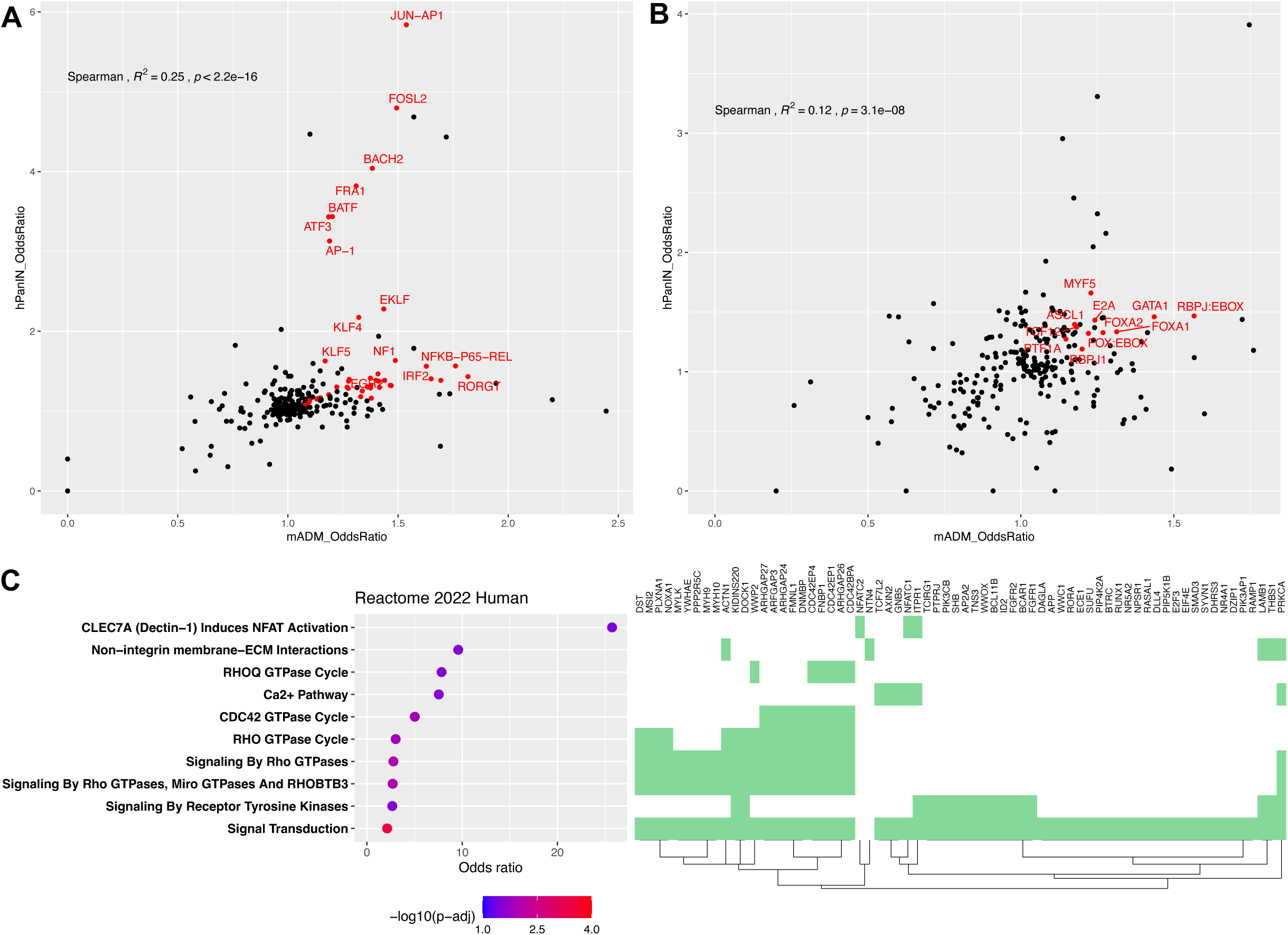
Consistent patterns among ADM DMRs and human PanIN DMRs. A-B. Correlation of motif enrichment analysis between mouse ADM and human PanIN. A. X-axis: motif enrichment odds ratio for mouse AK model D4 ADM vs acinar *hypomethylated* DMRs. Y-axis: motif enrichment odds ratio for human PanIN vs acinar *hypomethylated* DMRs. B. X-axis: motif enrichment odds ratio for mouse AK model D4 ADM vs acinar *hypermethylated* DMRs. Y-axis: motif enrichment odds ratio for human PanIN vs acinar *hypermethylated* DMRs. C. GSEA of shared DMR genes between mouse AK model D4 ADM vs acinar and human PanIN vs acinar (left), with genes contributing to each enriched gene set (right).

**Extended Data Figure 7.**
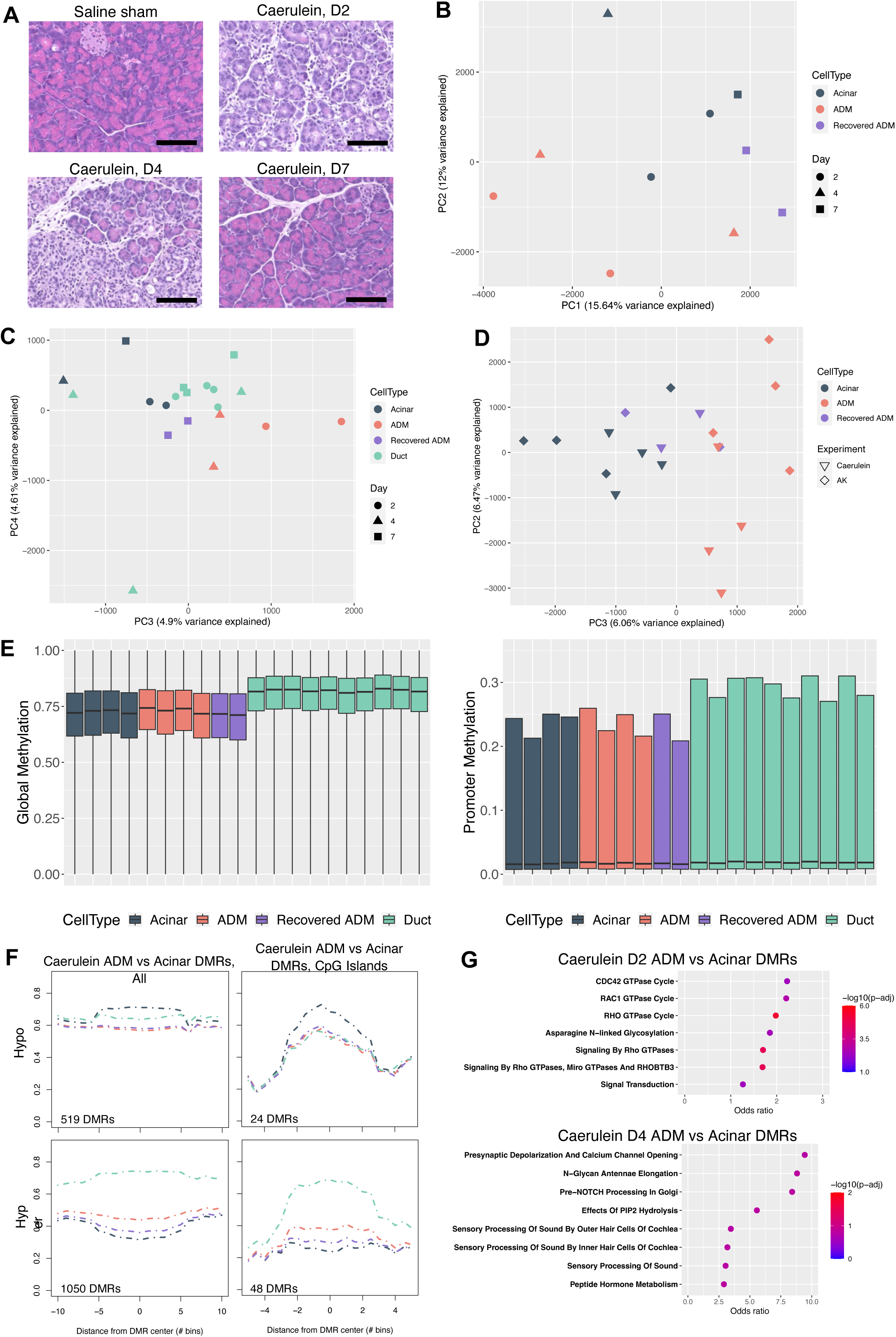
Methylation profiles in widely used caerulein model of pancreatitis. A. Hematoxylin and eosin (H&E) staining of normal control, and Day 2, Day 4, and Day 7 pancreata in caerulein model of acute pancreatitis. B. PCA of laser capture microdissection-purified acini, ADM lesions, and recovered ADM lesions at PCs 1 and 2. C. PCA of laser capture microdissection-purified acini, ADM lesions, recovered ADM lesions, and ducts at PCs 3 and 4. D. PCA of laser capture microdissection-purified acini, ADM lesions, recovered ADM lesions combined from both the AK and caerulein experiments at PCs 2 and 3. E. Boxplot of genome-wide (left) and promoter-specific (right) CpG methylation values for all exocrine samples. F. Meta-region plots of caerulein model ADM vs acinar DMRs summarized across all regions (left) and across CpG islands only (right) with 50% width buffer on each side. G. GSEA of genes overlapping caerulein model D2 ADM vs acinar DMRs (top) and D4 ADM vs acinar DMRs (bottom) using Reactome 2022 Human gene sets.

**Extended Data Figure 8.**
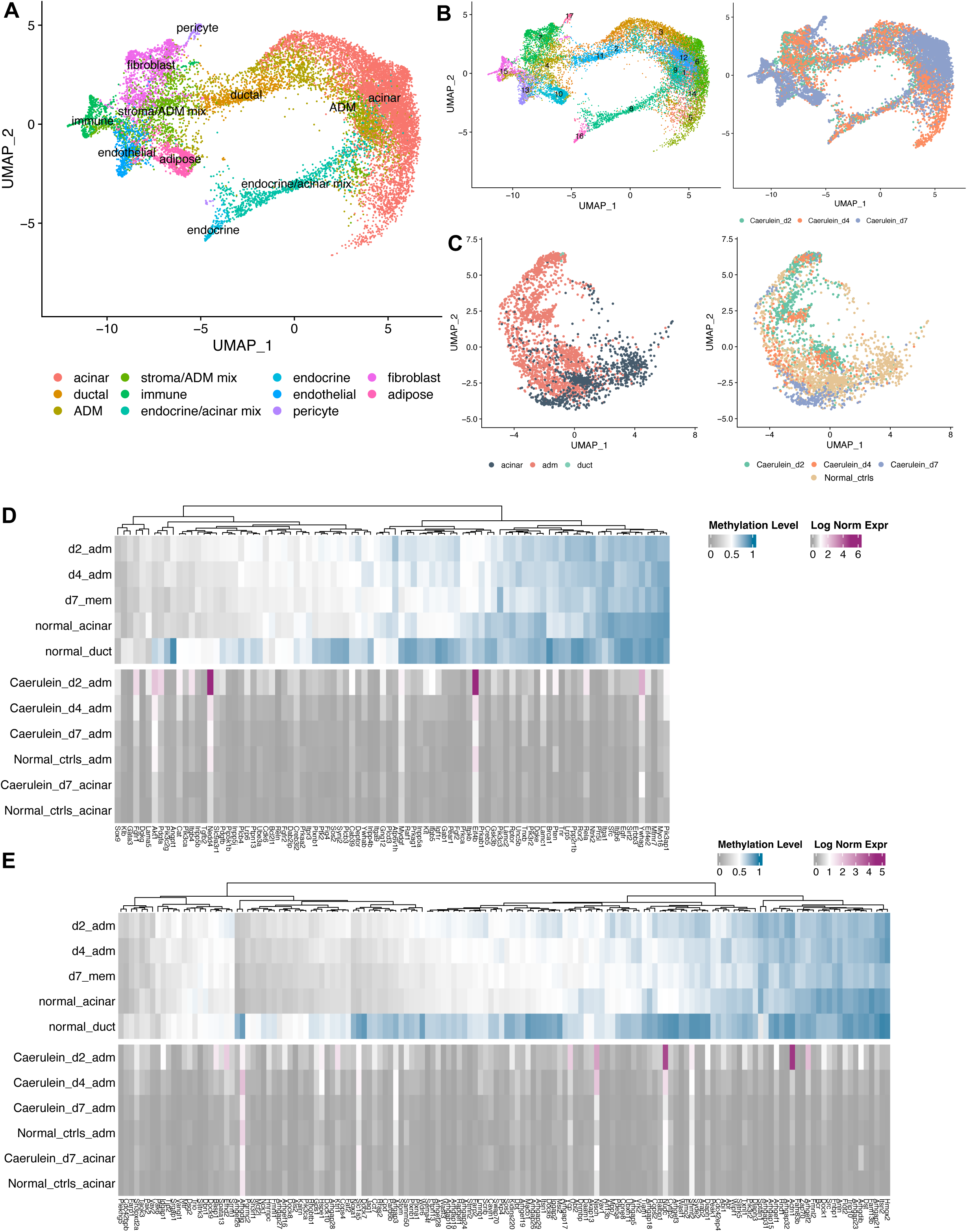
Gene expression patterns in caerulein model pancreata. A. UMAP projection of caerulein model Visium ST spots (n = 16,079 spots) by manually annotated major cell type. B. UMAP projection of ST spots by leiden clustering (left) and timepoint (right) for all caerulein model samples. C. UMAP projection of high-purity exocrine spots from caerulein model pancreata by deconvolution-assigned cell type (left) and by timepoint (right). D. Summary heatmap of methylation level and normalized expression level at Reactome- and KEGG-annotated Pi3k GTPase-related genes for each caerulein sample. E. Summary heatmap of methylation level and normalized expression level at Reactome- and KEGG-annotated Rho/Rac/Cdc42 GTPase-related genes for each caerulein sample. Normal acinar, n = 4 mice; day 2 ADM, n = 2 mice; day 4 ADM, n = 2 mice; day 7 ADM, n = 2 mice; duct, n = 10 mice.

**Extended Data Figure 9.**
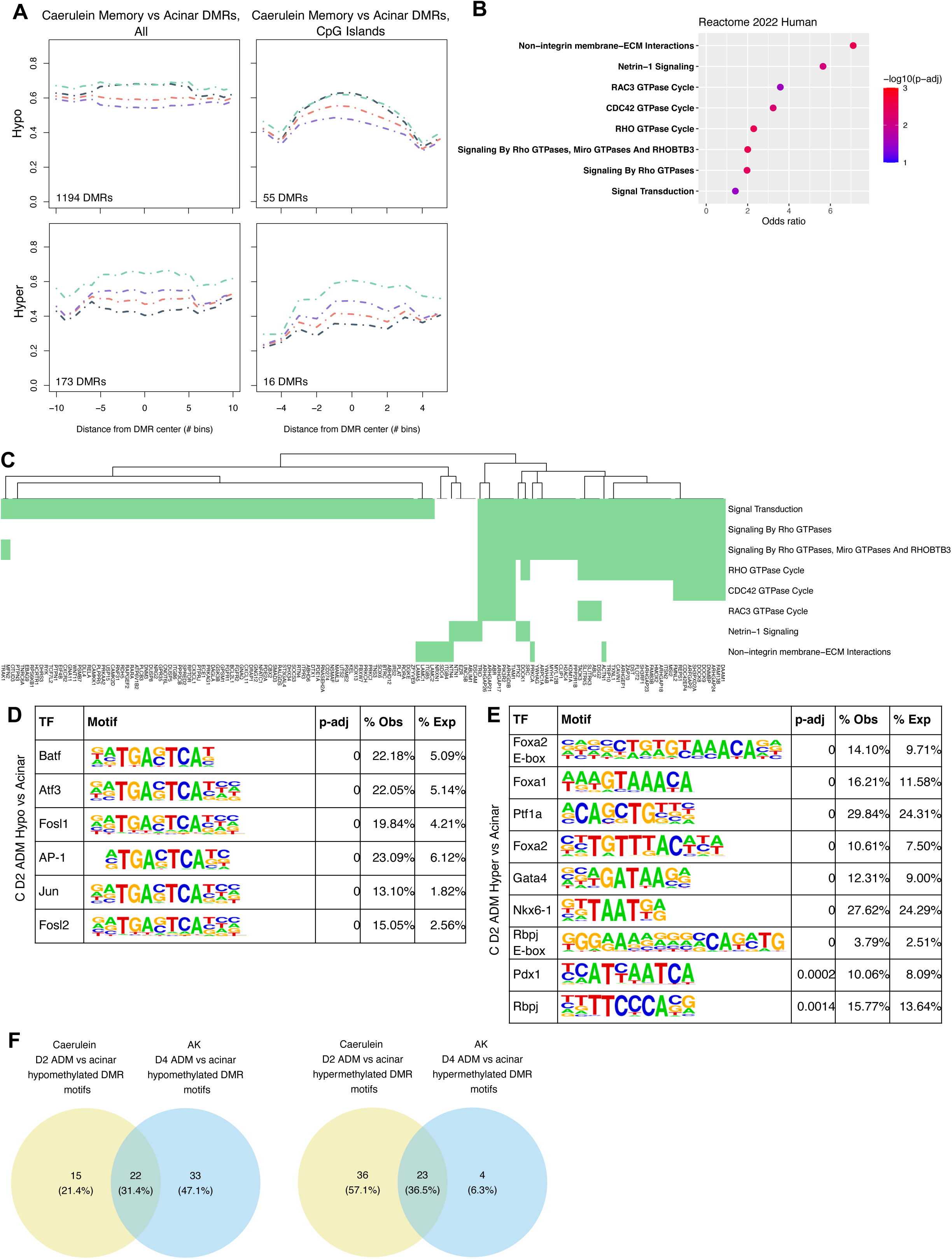
DNA methylation memory in caerulein model of ADM. A. Meta-region plots of caerulein model D7 vs acinar DMRs summarized across all regions (left) and across CpG islands only (right) with 50% width buffer on each side. Normal acinar, n = 4 mice; day 2 ADM, n = 2 mice; day 4 ADM, n = 2 mice; day 7 ADM, n = 2 mice; duct, n = 10 mice. B. GSEA of genes overlapping caerulein model D7 vs acinar DMRs using Reactome 2022 Human gene sets. C. Genes (columns) overlapping caerulein model D7 ADM vs normal acinar DMRs that contribute to each significantly enriched gene set (rows) as in B. D. Motif enrichment analysis results at caerulein ADM vs acinar hypo DMRs at Day 2. E. Motif enrichment analysis results at caerulein ADM vs acinar hyper DMRs at Day 2. F. Venn diagrams indicating the overlap of hypomethylated DMR motifs and hypermethylated DMR motifs between the Caerulein and AK models.

**Extended Data Figure 10.**
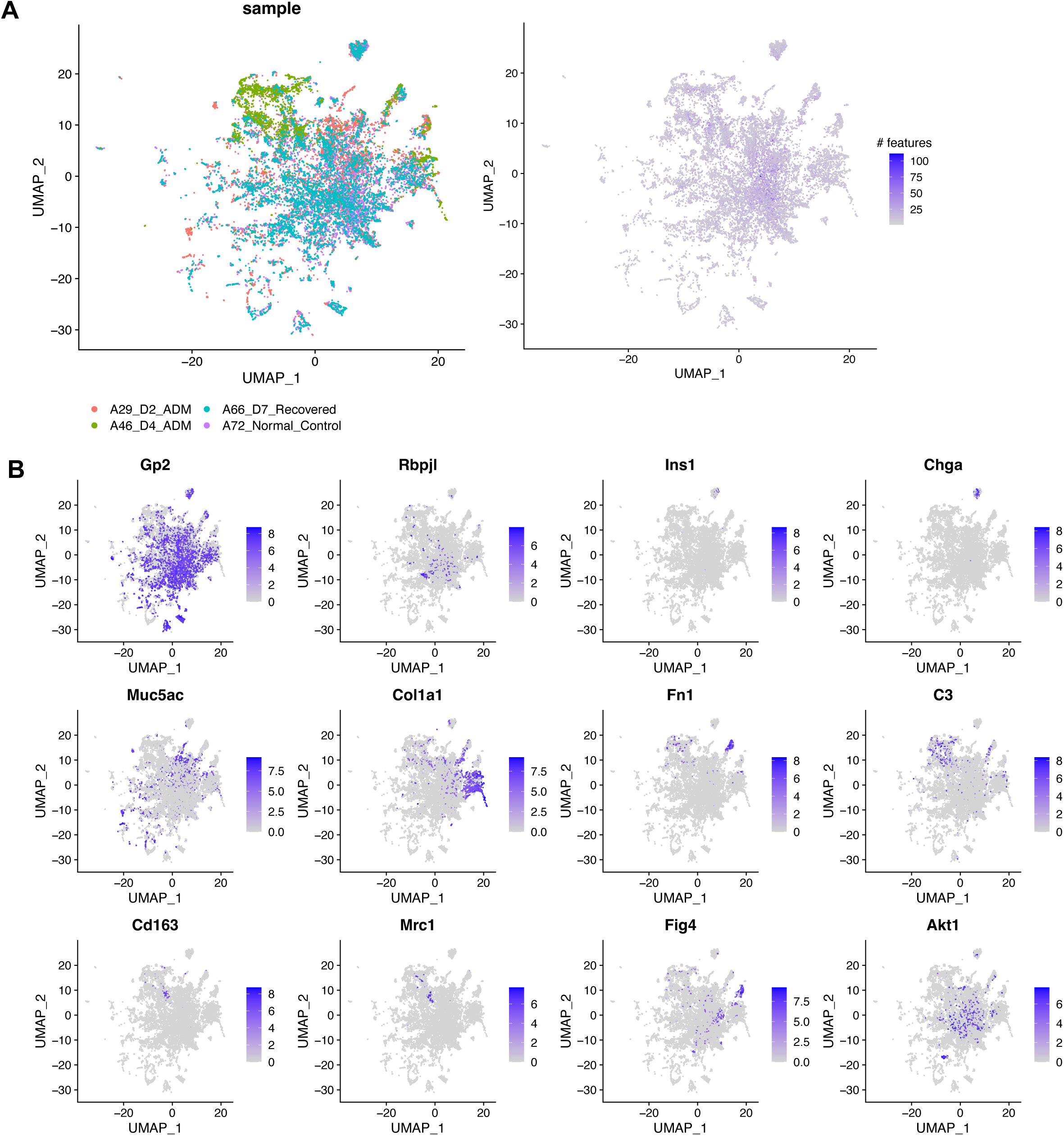
MERSCOPE targeted spatial transcriptomics on AK model pancreata. A. UMAP projection of all MERSCOPE cell profiles (n = 19776 cells) by timepoint (left) and by number of genes detected in each cell (right) B. Normalized expression of canonical marker genes for acini (*Gp2, Rbpjl*), islet cells (*Ins1, Chga*), PanINs (*Muc5ac*), fibroblasts (*Col1a1, Fn1*), macrophages (*Cd163, Mrc1*), and the Pi3k pathway (*Fig4, Akt1*).

## Supplementary Tables

**Supplementary Table 1.**

Mouse pancreas and body weights for AK and caerulein experiments.

**Supplementary Table 2.**

Summary of pancreatic samples and omics QC metrics.

**Supplementary Table 3.**

Genomic variants for all AK experiment mice as called by freebayes and maftools.

**Supplementary Table 4.**

Genes overlapping differentially methylated regions (DMRs) for:

- AK D2 ADM (n = 2 mice) vs normal acini (n = 4 mice)
- AK D4 ADM (n = 2 mice) vs normal acini (n = 4 mice)
- AK D7 (recovered, n = 2 mice) vs normal acini (n = 4 mice), all
- AK D7 (recovered, n = 2 mice) vs normal acini (n = 4 mice), where the D7 methylation level was more extreme than that of D2 and D4
- Caerulein D2 ADM (n = 2 mice) vs normal acini (n = 4 mice)
- Caerulein D4 ADM (n = 2 mice) vs normal acini (n = 4 mice)
- Caerulein D7 (recovered, n = 2 mice) vs normal acini (n = 4 mice)

Positive areaStat: hypermethylated in group 1.

**Supplementary Table 5.**

Publicly available single-cell mouse pancreas studies mined from NCBI Gene Expression Omnibus (GEO) used for deconvolution of spatial transcriptomics data.

**Supplementary Table 6.**

Homer motif analysis results for AK D4 and Caerulein D2 comparisons.

**Supplementary Table 7.**

MERFISH 500-gene target panel comprising a mixture of canonical marker genes, differentially methylated genes identified via WGBS, differentially expressed genes identified via 10X Visium ST, and TFs identified via differential motif enrichment analysis.

**Supplementary Table 8.**

qPCR primer sequences.

